# Combined landscape of single-nucleotide variants and copy-number alterations in clonal hematopoiesis

**DOI:** 10.1101/2021.03.05.433727

**Authors:** Ryunosuke Saiki, Yukihide Momozawa, Yasuhito Nannya, Masahiro M Nakagawa, Yotaro Ochi, Tetsuichi Yoshizato, Chikashi Terao, Yutaka Kuroda, Yuichi Shiraishi, Kenichi Chiba, Hiroko Tanaka, Atsushi Niida, Seiya Imoto, Koichi Matsuda, Takayuki Morisaki, Yoshinori Murakami, Yoichiro Kamatani, Shuichi Matsuda, Michiaki Kubo, Satoru Miyano, Hideki Makishima, Seishi Ogawa

**Author notes:** **Correspondence should be addressed to**: Seishi Ogawa. **Conflict of interest disclosure**: The authors declare no conflict of interest.

## Abstract

Implicated in the development of hematological malignancies (HM) and cardiovascular mortality, clonal hematopoiesis (CH) in apparently healthy individuals has been investigated by detecting either single-nucleotide variants and indels (SNVs/indels) or copy number alterations (CNAs), but not both. Here by combining targeted sequencing of 23 CH-related genes and array-based CNA detection of blood-derived DNA, we have delineated the landscape of CH-related SNVs/indels and CNAs in a general population of 11,234 individuals, including 672 with subsequent HM development. Both CH-related lesions significantly co-occurred, which combined, affected blood count, hypertension, and the mortality from HM and cardiovascular diseases depending on the total number of both lesions, highlighting the importance of detecting both lesions in the evaluation of CH.

## Introduction

The presence of clonal components in an apparently normal hematopoietic compartment, or clonal hematopoiesis (CH), has been drawing an increasing attention of recent years^1,2^. Although suggested only indirectly by skewed chromosome X inactivation in early studies^3-7^, CH has recently been demonstrated by detecting copy number alterations (CNAs) in the peripheral blood samples from large cohorts of individuals without blood cancers using single-nucleotide polymorphism (SNP) array data from genome-wide association studies (GWAS)^8-11^. Showing a substantial overlap to those characteristic of hematological malignancies (HM), CNAs were shown to be associated with an elevated risk of developing HM^8,9^. More recently, CH has also been detected by the presence of somatic single-nucleotide variants and indels (SNVs/indels) in the peripheral blood of apparently healthy individuals^**12-15**^ and cancer patients^**16**,**17**^ using next generation sequencing. In addition to its link to HM, CH as detected by SNVs/indels has been highlighted by its unexpected association with a significantly increased risk for cardiovascular diseases (CVD)^12,13,18,19^.

Regardless of the type of genetic lesions by which it is detected, CH is strongly age-related with an increasing frequency in the elderly^8-13^. With substantially improved technologies to identify CNAs and somatic SNVs/indels, a complete registry of CNAs and SNVs/indels associated with CH has been elucidated, which are thought to involve virtually every individual in the extreme elderly^20,21^. However, to date, no studies have evaluated both CNAs and SNVs/indels together at a comparable sensitivity in a large cohort of a general population, although they have recently been investigated in a cancer population, where many had been treated with chemo/radiotherapy^22^. What is the landscape of CH recognized by combining both CNAs and SNVs/indels in a general population? Are there any interactions between SNVs/indels and CNAs that shaped the landscape of CH? How are hematological phenotypes affected by both CH-related lesions? How does it affect HM and CVD risks? These are the key questions to be answered for better understanding of CH and its implication in HM and CVD.

In the present study, for the purpose of delineating the combined landscape of common driver SNVs/indels and CNAs in CH, we performed SNP array-based copy number analysis and targeted sequencing of major CH-related genes on blood-derived DNA from the Biobank Japan (BBJ)^23^, which had been SNP-typed for GWAS studies for common diseases, including hypertension, diabetes, autoimmune diseases and several solid cancers^23^. We then investigated the combined effect of both CH-related lesions on clinical phenotypes and outcomes, particularly that on the mortality from HM and CVD.

## Identification of CH-related SNVs/indels and CNAs

We enrolled a total of 11,234 subjects from the BBJ cohort (n=179,417), in which SNP array analysis of peripheral blood-derived DNA had been performed for large-scale GWAS studies for common diseases (Supplementary Table 1,2) (https://biobankjp.org/info/pdf/sample_collection.pdf)^23^. Among these 10,623 were randomly selected from 60,787 cases who were aged ≥60 years at the time of sample collection and were confirmed not to have solid cancers as of March 2013. This randomly selected set included 61 cases who were known to develop and/or die from HM as of March 2017. The remaining 611 consisted of all cases from the entire BBJ cohort who were confirmed to develop and/or die from HM as of the same date but were not included in the randomly selected 10,623 cases. In total, 672 cases were reported to have HM in the entire BBJ cohort, which included 215 myeloid, 420 lymphoid, and 37 lineage-unknown tumors (Extended Data Fig. 1a). For these 11,234 cases, SNVs/indels in blood were investigated using multiplex PCR-based amplification of exons of 23 CH-related genes, followed by high-throughput sequencing (Online methods).^24^ Sensitivity of SNV detection according to *in silico* simulations using known SNPs was >94% for 3% variant allele frequency (VAF) and >74% for 2% VAF, but <20% for 1% VAF with a mean depth of ∼800x (Supplementary Fig. 1a-b).

**Fig. 1.**
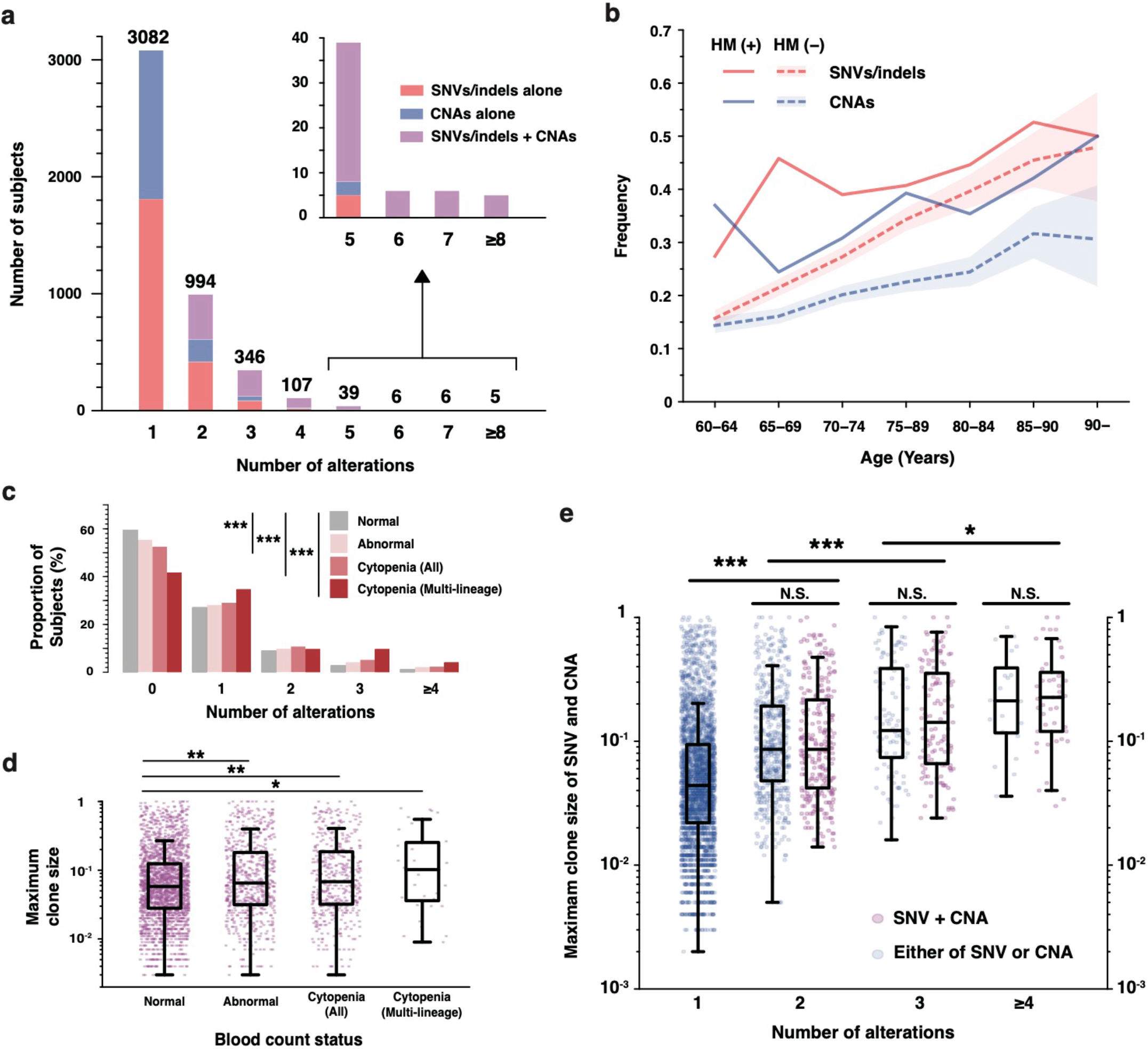
Landscape of SNVs/indels and CNAs in clonal hematopoiesis. a, Distribution of the number of genetic alterations in each subject. Subjects with SNV/indels alone, with CNAs alone, or with both of them are illustrated by different colors. b, The prevalence of CH-related SNVs/indels and CNAs in subjects with or without HM events, according to age. Colored bands represent the 95% confidence intervals. c, Number of coocurring alterations in those with subjects with abnormalities in blood cell counts, or cytopenia. d, Maximum cell fraction of CH-related alterations in subjects with abnormalities in blood cell counts, and in those with cytopenia. e, Dotplot of maximum cell fractions of SNVs/indels or CNAs across different numbers of coocurring alterations. Cell fractions of SNVs/indels are defined as 2 times VAF. Those with both of SNVs/indels and CNAs are shown in purple, while those with either are shown in blue. In panel (d,e), unclassifiable CNAs were excluded because we cannot calculate their precise cell fractions. The box plots indicate the median, first and third quartiles (Q1 and Q3) and whiskers extend to the furthest value between Q1 1.5-xthe interquartile range (IQR) and Q3 + 1.5xIQR. *P* values were calculated by wilcoxon rank-sum test. N.S., not significant;, * *P < 0*.*05;* **, *P*<0.001; ***, *P*<0.0001.

In total, we called 4,056 SNVs/indels (2,750 SNVs and 1,306 indels) in 3,071 (27.3 %) subjects, of which 2,312 (20.6%) had one, 586 (5.2%) two, and 173 (1.5%) ≥3 SNV/indels (Fig. 1a). Their VAFs widely distributed from 0.5% to 85.6% with a median of 3.0% (Supplementary Fig. 1c). Age-dependence of CH-related SNVs/indels was evident (Fig. 1b). In accordance with previous reports, *DNMT3A* (13.5%), *TET2* (9.5%), *ASXL1* (2.2%), and *PPM1D* (1.4%) were most frequently mutated (Extended Data Fig. 2a,c). Several combinations of genes, including *TET2/DNMT3A, ASXL1*/*TET2, ASXL1*/*CBL, SRSF2*/*TET2*, and *SRSF2*/*ASXL1*, were more frequently co-mutated than expected only by chance (OR: 1.53-6.53, *q*<0.05) (Extended Data Fig. 2d). Of interest, many of these combinations are also co-mutated in myeloid neoplasms with large VAF values^25-27^, suggesting the presence of these combinations of SNVs/indels in the same cell fraction. This was also expected for some cases having a large (>50%) sum of VAFs of relevant SNVs/indels (“pigeonhole principle”),^28^ although it was not determined whether or not these combinations of SNVs/indels affected the same cell populations in the vast majority of cases (Extended Data Fig. 2e-i).

**Fig. 2.**
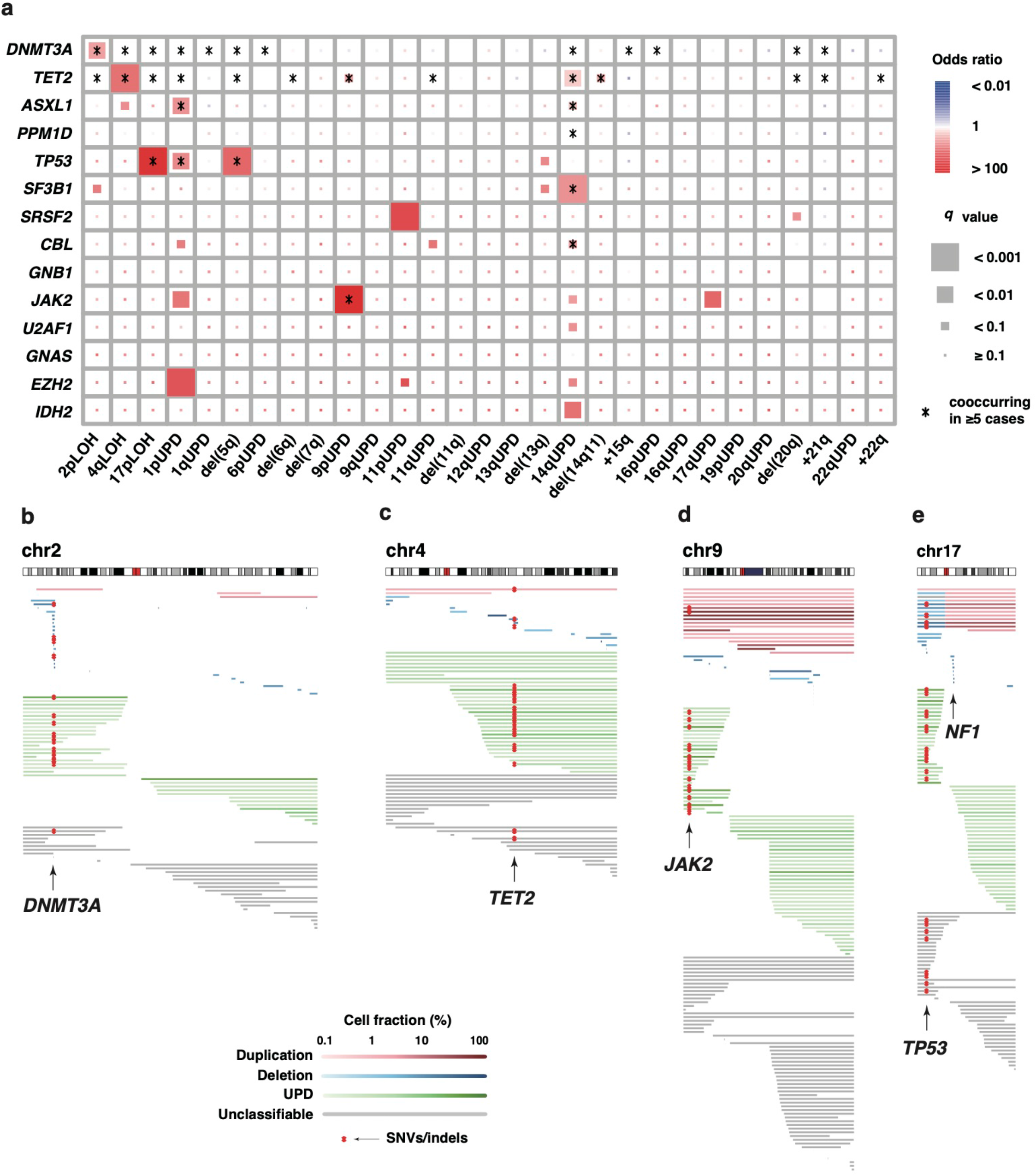
Cooccurrences of SNVs/indels and CNAs in clonal hematopoiesis. a, The correlations between individual SNVs/indels and CNAs. The size of rectangles indicate the significance of correlations. Red rectangles represent positive correlations while blue rectangles represent negative correlations. Combinations of SNVs/indels and CNAs seen in 5 or more subjects are indicated by asterisks. b-e, The distributions of CNAs on chromosome 2 (b), 4 (c), 9 (d), and 17 (e). Horizontal bars represent CNAs, and cooccurring SNVs/indels in *DNMT3A, TET2, JAK2*, and *TP53* are indicated by red asterisks. Colors of horizontal bars represent the types and cell fractions of CNAs. Allele imbalances which cannot be classified into any of UPD, deletion, or duplication are indicated as unclassifiable CNAs (gray).

CNAs data were available from the previous study^21^, in which SNP array-based copy number detection in blood-derived DNA was performed for a larger cohort of BBJ cases (n=179,417), including all the cases enrolled in the current study (n=11,234). In total, 2,797 CNA-positive regions/segments were identified in 2,254 (20.1%) cases (Extended Data Fig. 3, Online methods), of which 413 (3.7%) had multiple CNAs (Fig. 1a). Reflecting a higher age distribution of the current cohort, the frequency of CNAs was higher than that in the entire BBJ cohort^21^, even though age-stratified frequencies were almost equivalent between both cohorts (Fig. 1b). Estimated mutant cell fractions (MCF) for CNAs were ranged from 0.2% to 93.2% with a median of 2.0% with FDR<0.05, where a substantial number (n=461) of CNAs were seen in a cell fraction of ≤1%, which was below the limit of detection for SNVs/indels. Thus, smaller clones were detected through CNAs, particularly copy-neutral loss-of-heterozygosity (CN-LOH) or uniparental-disomy (UPD), compared with through SNVs/indels (Supplementary Fig. 1c).

**Fig 3.**
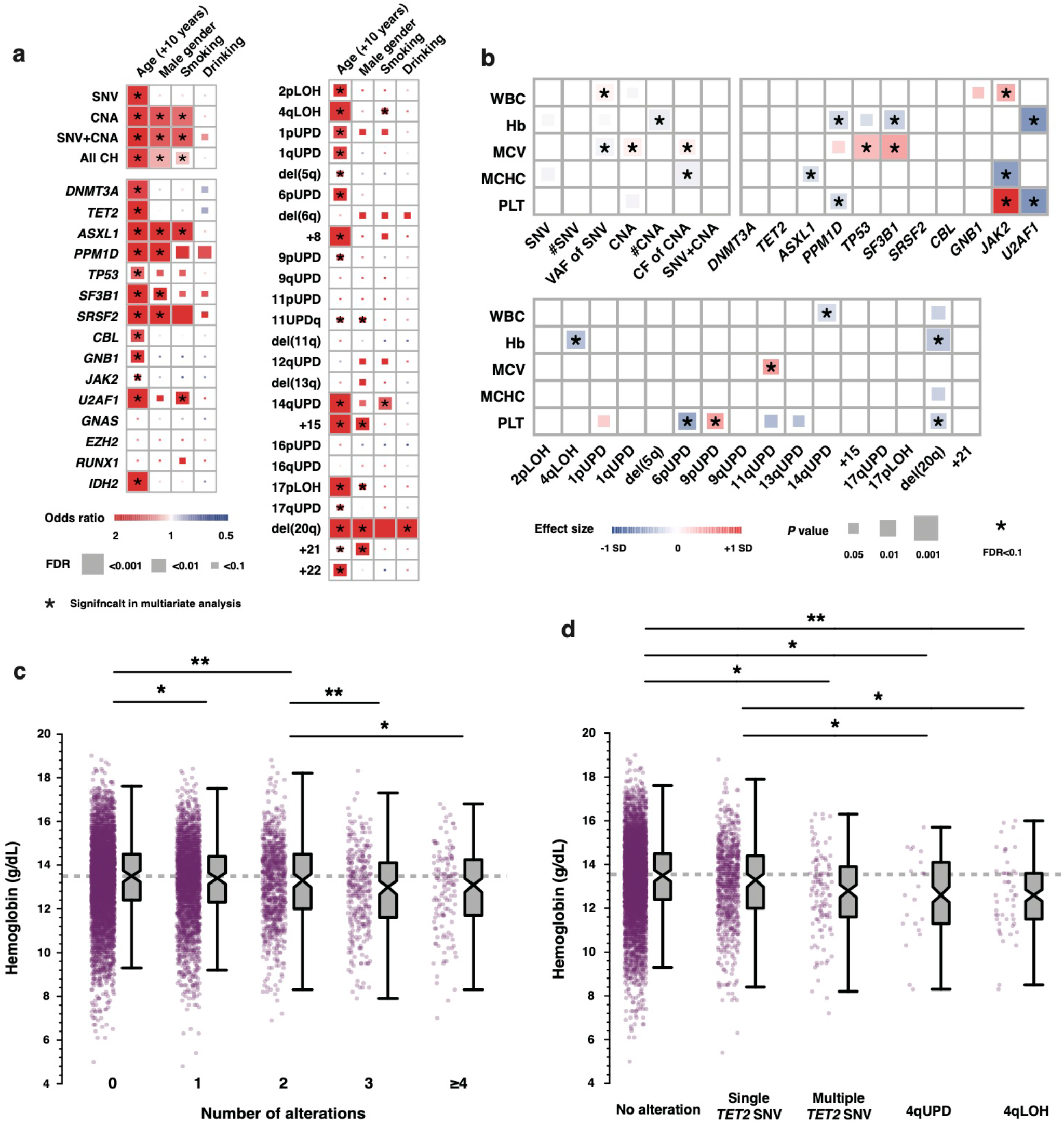
Risk factors for CH and effects on blood counts. a, Correlations of genetic alterations with age, male gender, history of smoking and drinking. Sizes and colors of rectangles represent the significance and effect size, respectively. Asterisks indicate the clinical factors significantly correlated with each alteration in multivariate logistic regression (*P*<0.05). c, Correlations between genetic alterations and blood counts. The sizes and colors of rectangles indicate the significance, and effect size of correlations. Correlations significant after correction for multiple testing (FDR<0.1) are indicated by asterisks. WBC: white blood cell, Hb: hemoglobin, MCV: mean corpuscular volume, MCHC: mean corpuscular hemoglobin concentration, Plt: Platelet. c, Distributions of hemoglobin in subjects with different number of alterations. d, Distributions of hemoglobin in subjects with no alterations, with single SNV/indel in *TET2* (Single *TET2* SNV), multiple SNVs/indels (Multiple *TET2* SNV) in *TET2*, with 4qUPD, or with any allelic imbalances in 4q (4qAI) are illustrated in dot plots and boxplots. In all box plots, the median, first and third quartiles (Q1 and Q3) are indicated and whiskers extend to the furthest value between Q1 -1.5xthe interquartile range (IQR) and Q3 + 1.5xIQR. \**p* < 0.05; ** *p* < 0.01.

We found 27 significantly recurrent CNAs, many of which are also commonly seen in HM, supporting a pathogenic link between CH and leukemogenesis (Extended Data Fig. 4a-c). In accordance with previous reports^8-11^, 14qUPD, +21q, del(20q), and +15q were among the most frequent CNA lesions (Extended Data Fig. 2b,c), while del(20q), 16pUPD, and 17pUPD showed the largest mean clone size (Supplementary Fig. 2). Several CNAs, such as 14qUPD and +21, showed higher frequencies than reported in western populations, which is likely due to a higher sensitivity for detecting CNAs in this study compared with that in previous studies in western populations^8-11^; when confined to lesions with ≥5% cell fractions, the difference across studies becomes less conspicuous for many CNA targets (Extended Data Fig. 4d,e). Nevertheless, even considering the different sensitivities, several CNAs, including +15, del(14q), del (9q), del(20q) and del(13q), still showed a different frequency across studies in both populations ^21^, suggesting an ethnic difference in positive selection of CH-related CNAs (Extended Data Fig. 4e), although the exact genetic basis of the ethnic difference is largely unclear for most CNAs.

**Fig. 4.**
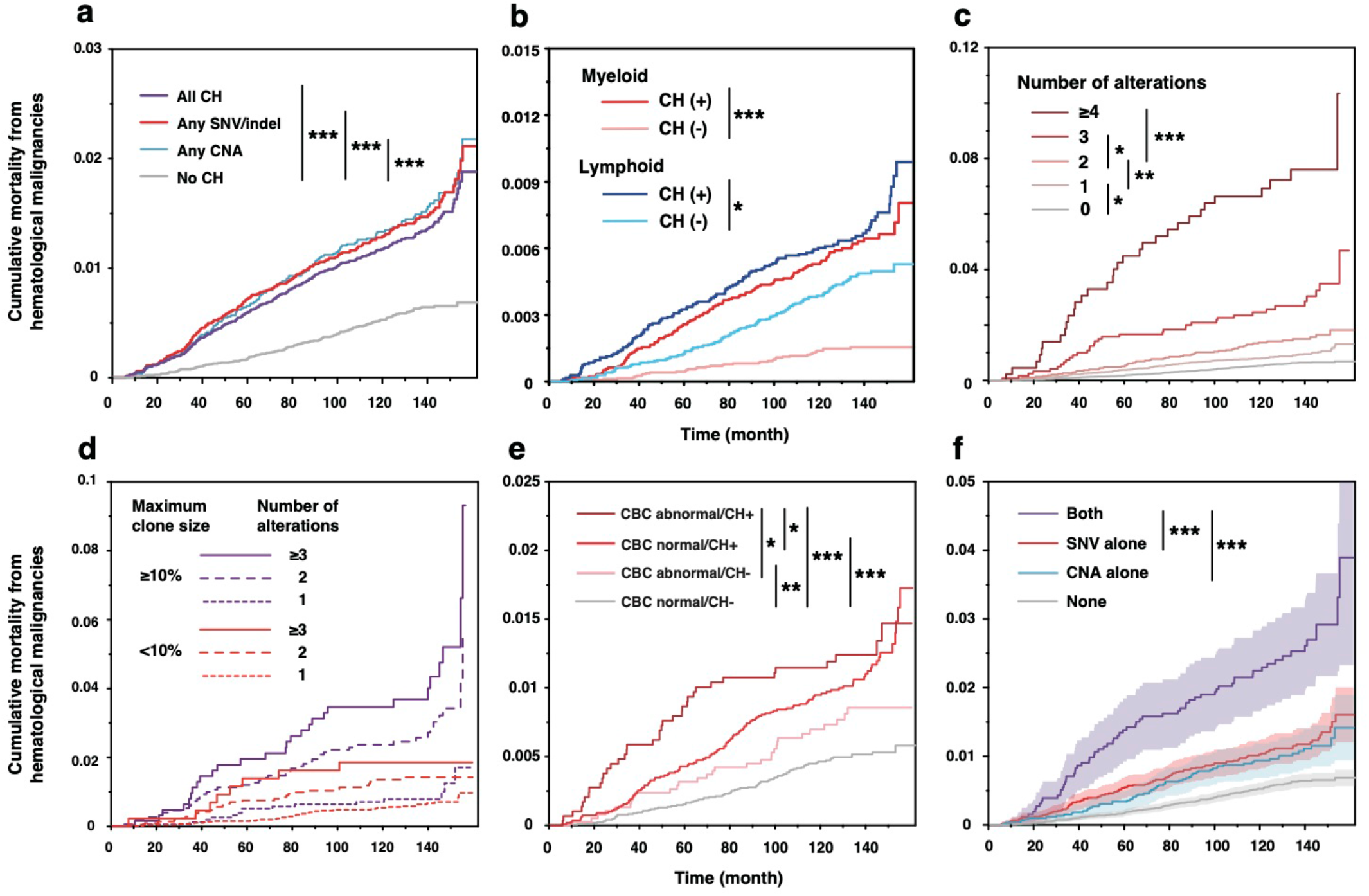
Impact of CH on mortality from hematological malignancies. a, Cumulative mortality from HM in subjects with any CH, any SNV/indel, any CNA, or without CH are shown. b, Cumulative mortality from myeloid and lymphoid malignancies in subjects with or without CH are shown. c, Cumulative mortality from HM in subjects with different numbers of CH-related alterations. d, Cumulative mortaliry from HM in subjects with different numbers of coocurring alterations and maximum clone sizes (<10% or .210%). Cell fractions of unclassifiable CNAs were regarded to be smaller than 10%. e, Cumulative mortality from HM in subjects with or without abnormalities in complete blood counts (CBC) and/or CH. f, Cumulative mortality from HM in subjects with both SNV/indels and CNA, SNV/indels alone, CNAs alone, and without any alterations are shown. Colored bands indicarte 95% confidence intervals. \**P* < 0.05, \*\**P* < 0.001, \*\*\**P* < 0.0001.

## Combined landscape of SNVs/indels and CNAs

When SNVs/indels and CNAs were combined, CH was demonstrated in 4,242 (40%) of randomly selected 10,623 cases who were ≥60 years of age with no reported cancer history and in 376 (56%) of 672 cases who developed HM, where 38 of the 376 were <60 years old. Combining both lesions, more subjects (n=1,503) had two or more lesions than judged by SNVs/indels (n=759) or CNA alone (n=413) (Fig. 1a). The frequency of CH and the total number of CH-related lesions, as well as the maximum estimate of clone size in CH(+) cases, were significantly larger in individuals with abnormal blood counts, particularly those with cytopenias, compared with those with completely normal blood counts, depending on the number of blood lineages involved (Fig. 1c,d). A similar landscape of combined CH-lesions was observed in an independent cohort of 8,023 solid cancer patients from The Cancer Genome Atlas (TCGA), although the sensitivity of CH-lesions, particularly CNAs, was substantially lower than the current study due to a lower coverage of exome sequencing and a less accurate haplotype phasing required for sensitive CNA detection (Extended Data Fig. 5a,b,c).

**Fig 5.**
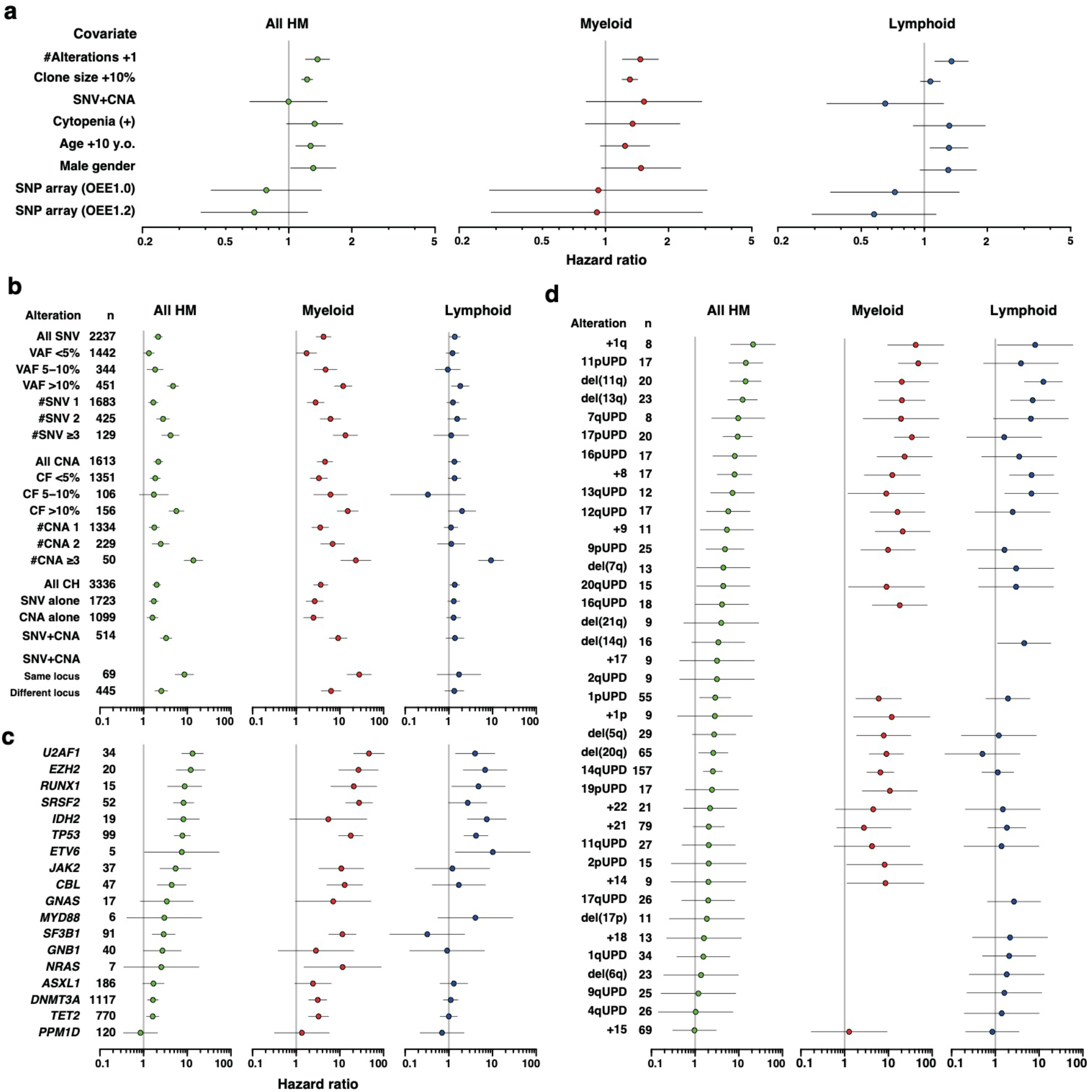
Impact of CH-related alterations on mortality from HM. a-d, Hazard ratios for mortality from All hematological malignancies (All HM), myeloid neoplasms, and lymphoid neoplasms. Error bars indicate 95% confidence intervals. In panel (a), hazard ratios of the indicated covariates calculated by multivariate Fine-Gray regression are shown. In panel (b-d), hazard ratios of the indicated alterations are calculted in comparison with subjects without any alteration. Hazard ratios are not shown for alterations without any event. Cell fractions of unclassifiable CNAs were regarded to be zero in panel (a), and smaller than 5% in panel (b). n, number of cases with the indicated alterations; #Alteration, additional one alteration; Clone size +10%, 10% increase in cell fraction; SNV+CNA, coocurrence of both SNVs/indels and CNAs; #SNV, number of SNVs/indels; CF, cell fraction of CNAs; #CNA, number of CNAs.

Accounting for 7% of the total cohort and 16% of all CH(+) cases, 740 individuals harbored both types of lesions, which were significantly more frequent than expected only by chance (Extended Data Fig. 2j), even after their age was adjusted (odds ratio [OR]=1.3; *P*=0.0003, age-stratified permutation test) (Online methods). SNVs/indels in *TP53, TET2, JAK2, SF3B1*, and *U2AF1*, and less significantly in *DNMT3A, CBL*, and *SRSF2*, was accompanied by significantly more CNAs (Supplementary Fig. 3). The number of cases with multiple CH-related lesions was also significantly larger than expected from the number of all CH-related lesions (*P*=0.0067). The significantly higher frequency of cases with both SNVs/indels and CNAs (*P*<0.0001) and those with multiple lesions (*P*<0.0001) were confirmed in the TCGA cohort. These observations raise a possibility that it might be the total number of lesions, rather than the combination of SNVs/indels and CNAs, that is relevant to the positive selection in CH, in which multiple CH-related lesions in the same cell contributed to positive selection in a substantial number of cases with multiple CH-lesions. In support of this, the maximum clone size in CH(+) cases significantly correlated with the total number of CH-related SNVs/indels and CNAs, but not their combinations per se (Fig. 1e).

Co-occurring multiple lesions were judged to be present in the same cell in 73 cases on the basis of their large (>1.0) clone size sum^28^, of which 8 were combinations between SNVs/indels and CNAs (Extended Data Fig. 2k). In the vast majority of cases, we could not determine the cellular compartment of multiple lesions due to small clone size of both lesions, which would be better addressed using single cell-based sequencing. A representative case was shown in Supplementary Fig. 4, in which the presence of both del(13q) and a *TET2*-involving SNV in the same cell compartment of myeloid lineages was demonstrated using single-cell sequencing (Supplementary Fig. 4a-d). Some combinations of SNVs/indels and CNAs were significantly more frequently observed than expected only by chance (Fig. 2a). Of particular interest among these were co-occurring SNVs/indels and CNAs affecting the same gene/locus. Overall, we found 88 cases having co-occurring SNVs/indels and CNAs affecting 8 genes/loci (Extended Data Fig. 6a), of which most frequently involved were *TP53* (with 17pLOH) (n=24, OR=60.6, *q*<0.001), *TET2* (with 4qLOH) (n=22, OR=10.8, *q*<0.001), *JAK2* (with 9pLOH/gain) (n=18, OR: 414, *q*<0.001), and *DNMT3A* (with 2pLOH) (n=16, OR=4.02, *q*=0.001), which were also found in the TCGA cases (Fig. 2a-e, Extended Data Fig. 5d). In reality, more cases are expected to have these combinations, because there were many ‘isolated’ LOH lesions or allelic imbalances affecting these loci that lacked accompanying SNVs/indels (n=64) (Fig. 2b-e), which were thought to escape from detection due to lower sensitivity of detecting SNVs/indels than CNAs (Supplementary Fig. 1a,c). In fact, using highly sensitive ddPCR assay targeting mutational hotspots, SNVs in *JAK2* and *TP53* were confirmed in 8 out of 48 and 24 out of 41 samples with isolated LOH at 9p and 17p, respectively (Supplementary Fig. 5). Representing well-known mechanisms of biallelic alterations of the relevant driver genes in myeloid malignancies, these combinations of lesions in CH are predicted to affect the same cell, being involved even in very early stages of positive selection in myeloid leukemogenesis^29-31^. SNVs/indels were most frequently associated with LOH when they affected *TP53* and *JAK2* in both myeloid malignancies^32,33^ and CH (Extended Data Fig. 6b), also supporting their role in the mechanism of biallelic alterations. Unfortunately, none of these cases satisfied the pigeonhole principle or no samples were available for single cell-sequencing analysis to directly confirm this at a single cell level. However, in the case of SNVs/indels associated with UPD, their presence in the same cell compartments in many cases was supported by a highly skewed distribution of mutant cell fractions of both lesions (Supplementary Fig. 6, Online methods).

**Fig 6.**
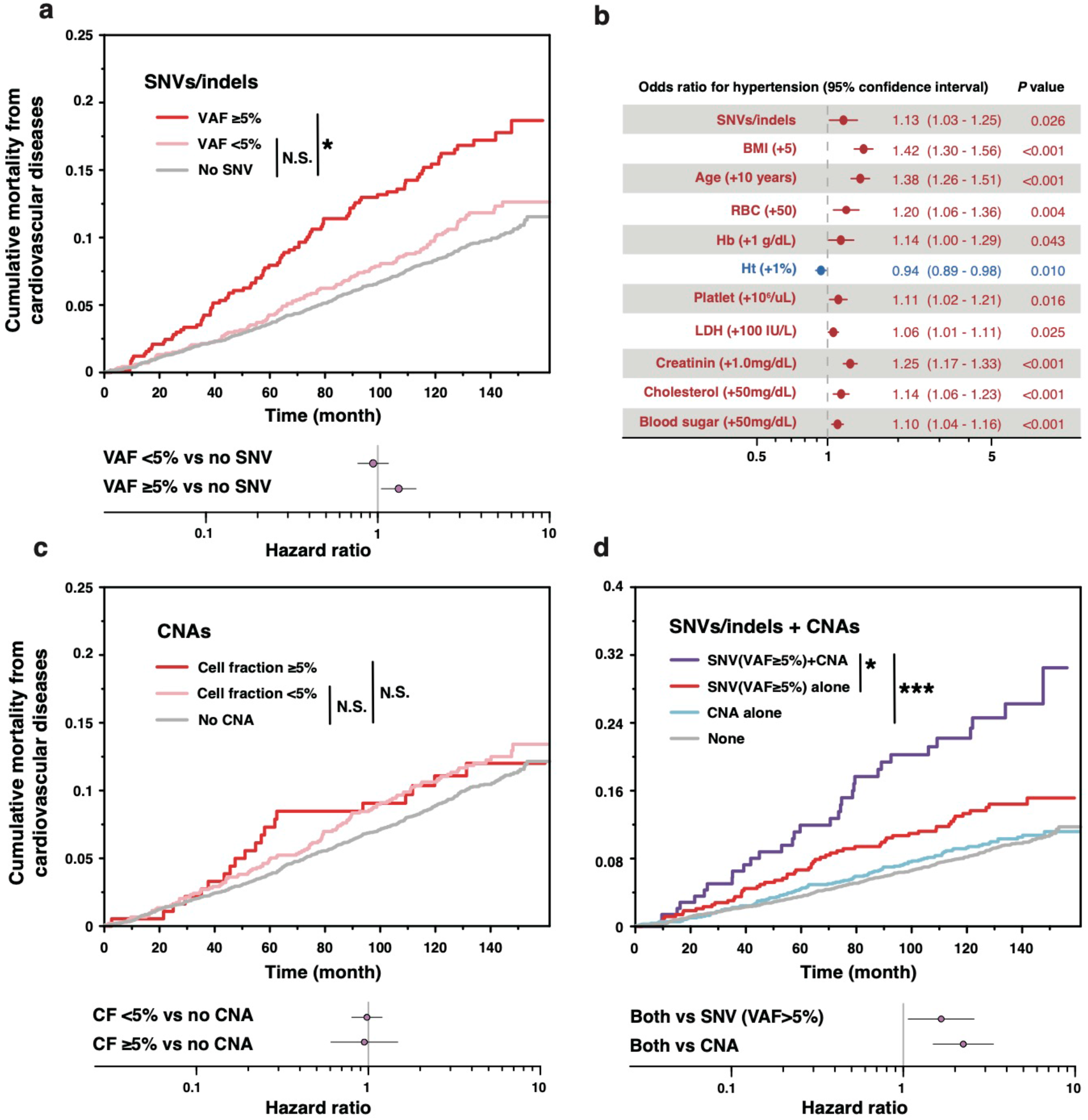
Effect of SNV/indels and CNAs on cardiovascular mortality. a, Cardiovascular mortality in subjects with SNV/indels (VAF ≥5% or <5%). and those without SNV/indels. Hazard ratios and P values are calculated in comparison with those without SNV/indels. b, Results of multivariate logistic regressions for the presence of hypertension. Explanatory variables were selected by stepwise method from following factors: presence of SNV/indels, CNAs, age (+10 years), male gender, BMI (+5), history of drinking and smoking, presence of diabetes mellitus, hyperlipid emia, hypertension, and 13 blood test values available in ≥70% of the subjects. Only remaining variables are shown. c, Cardiovascular mortality in subjects with CNAs (cell fraction ≥5% or <5%), and those without CNAs. Hazard ratios and P values for subjects are calculated in comparison with those without CNAs. d, Cardiovascular mortality in subjects with both SNV/indels (VAF≥5%) and CNAs (purple), with SNV/indels (VAF≥5%) alone (red), with CNAs alone (blue), and without SNV/indels (VAF≥ 5%) or CNAs (gray). Hazard ratios and *P* values were calculated by comparing those with both SNV/indels (VAF≥5%) and CNAs with those with SNV/indels (VAF≥5%) alone, or CNAs alone. In (a), (b) and (d), all comparisons were performed in multivariate models including age, gender, body-mass index, comorbidities (diabetes mellitus, hypertension, and dyslipidemia), history of smoking/drinking, and the versions of SNP array.

Besides SNVs/indels and CNAs affecting the same gene/locus, we also detected a significant combination between SNVs/indels in *TET2* and microdeletions of the *TCRA* (14q11.2 involving the) locus (n=7, OR=3.53, *q*=0.059), of which one case was reported to develop T-cell lymphoma (Fig. 2a and Extended Data Fig. 2l). This combination is of potential interest, given that *TET2* is frequently mutated in mature T-cell lymphomas^34^, particularly in follicular-helper T-cell-derived lymphomas, such as angio-immunoblastic T-cell lymphoma (AITL), which are also seen in *Tet2* knockdown mice^**35**^. Other potentially relevant combinations included *SF3B1*/14qUPD, *TET2*/14qUPD, *ASXL1*/1pUPD, *TP53*/1pUPD, and *TP53*/del(5q) (Fig. 2a), whose biological significance, however, is largely unclear except for the interplay between del(5q) and mutated-*TP53* intensively studied in myelodysplastic syndromes (MDS)^36,37^.

### Clinical associations with CH

Next, we investigated common demographic factors that may influence CH-related SNVs/indels and CNAs and the effect of both CH lesions on clinical features and outcomes. In addition to the large effect of age, several factors impacted on CNAs and/or SNVs/indels were observed. Male gender and smoking were significantly associated with SNVs/indels in *ASXL1, PPM1D*, splicing factors, and *TP53*, and with CNAs, particularly +15, del(20q), and +21 (with male gender), and 14qUPD (with smoking), many of which remained significant in multivariate analysis (Fig. 3a). The effect of alcohol consumption was less prominent and mostly confined to an increased incidence of del(20q). Although none of the subjects in our cohort had been diagnosed with HM at the time of sample collection, 1,314 cases had varying degrees of abnormal blood counts (Supplementary Table 3). Even though the landscape of CH in these cytopenic individuals at a glance was largely similar to that in non-cytopenic individuals (Extended Data Fig. 7a), cytopenic cases exhibited a significantly higher frequency of CH, where the frequency significantly correlated with the severity of cytopenia (Fig. 1c). In particular, individuals with abnormally high platelet counts had a higher frequency of *JAK2*-involving SNVs/indels and 9pUPD (OR=50.5, *q*<0.001 and OR=26.0, *q*=0.0017, respectively), while *U2AF1*-involving SNVs/indels, and del(20q) were more common in those with cytopenia of any sort (OR=7.39, *q*<0.001, and OR=3.10, *q*=0.015, respectively) (Extended Data Fig. 7b). Individuals with CH-related SNVs/indels had a higher frequency of cytopenia and exhibited lower hemoglobin and mean corpuscular hemoglobin concentration (MCHC), while CNAs was associated with lower WBC and platelet counts and larger mean corpuscular volume (MCV) (Fig. 3b). The number of all co-occurring alterations, SNVs/indels or CNAs, and VAF of SNVs/indels predicted significantly lower hemoglobin values, while MCF of CNAs predicted larger MCV and lower MCHC values (Fig. 3b,c, Extended Data Fig. 7c). As for individual alterations, SNVs/indels in *JAK2* were significantly correlated with high platelet counts (*q*<0.001) even when the analysis was confined to the individuals with normal blood counts. Moreover, we found significant associations of lower hemoglobin values with SNVs/indels in *TP53, PPM1D, SF3B1*, and *U2AF1*, and 4qUPD and del(20q), while SNVs/indels in *PPM1D, U2AF1*, 6pUPD, and del(20q) were associated with lower platelet counts and SNVs/indels in *TP53*, and *SF3B1*, and 11qUPD correlated with larger MCV (Fig. 3b, Extended Data Fig. 7c). VAF or cell fractions of SNVs/indels and CNAs were also predictive of the changes in hemoglobin, platelet counts, or MCV (Fig. 3b, Extended Data Fig. 7d). SNVs/indels in *TET2* alone were not associated with a reduced hemoglobin value (Fig. 3b). However, interestingly, we observed a significant association of lower hemoglobin values with multiple SNVs/indels in *TET2* and any allelic imbalance affecting 4q, which is most likely attributable to biallelic *TET2* alterations (Fig. 3b,d). We also tested the relationships between CH and values of other blood tests to reveal a negative correlation between *GNB1*-involving SNVs and uric acid achieved FDR<0.1 (Supplementary Fig. 7).

### Effect of SNVs/indels and CNAs on HM mortality

Among the major interests in the current study is the effect of SNVs/indels and CNAs on the risk of HM, particularly the combined effect of both CH-related lesions. To see this, we investigated the effect of CH on the cumulative mortality from HM using the Fine and Gray regression modeling in a case-cohort design^38^, where 7,937 of the 10,623 cases were regarded as a subcohort that were randomly selected from 43,662 cases who had been followed up for survival and cause of deaths on the basis of the vital statistics of Japan^39^ (Extended Data Fig. 1b). The median follow-up of these cases was 10.4 years (range, 0.01-13.5), during which 401 HM deaths were confirmed (Extended Data Fig. 1b). Age, sex, and versions of SNP array were adjusted and deaths from any causes other than HM were analyzed as competing risks.

In accordance with previous reports^22^, both SNVs/indels and CNAs were significantly associated with a higher mortality from HM than observed in CH(−) cases with an estimated cumulative 10-year mortality of 1.28% and 1.32%, respectively (Fig. 4a). Although lymphoid neoplasms accounted for two-thirds of all HM mortality in the cohort of 43,662 elderly cases, attributable mortality in CH(+) vs. CH(−) cases was ∼two times higher from myeloid neoplasms (0.39%) than lymphoid (0.21%) neoplasms (Fig. 4b) and the hazard ratio between CH(+) and CH(−) cases was >2.5 times larger for myeloid (3.64) than lymphoid (1.36) neoplasms (Fig. 5b). This suggests the predominant effects of CH on myeloid neoplasms, which is in line with the fact that most CH-related lesions targeted driver genes in myeloid neoplasms. The number of SNVs/indels and CNAs and the total number of CH-related lesions all significantly correlated with higher HM mortality (Fig. 4c, Extended Data Fig. 8a,b). While the maximum clone size of CH-related lesions correlated with the number of CH-related lesions (Fig. 1e), the former was also significantly associated with a higher HM mortality independently of the latter (Fig. 4d, Fig. 5a), which was in line with a previous observation that SNVs/indels correlated with development of HM only when they exhibited sufficiently large VAFs^40^. In univariate analysis, the largest risk of HM mortality was conferred by SNVs/indels of *U2AF1, EZH2, RUNX1, SRSF2* and *TP53*, +1q^11,14,21^ (Fig. 5c-d, Supplementary Fig. 8,9). As expected from a ∼2 times larger attributable mortality for myeloid than lymphoid malignancy, HRs and ORs were higher in myeloid than lymphoid HM for most of the lesions, with an exception of trisomy 12, which was associated with lymphoid, but not myeloid, neoplasms (Extended Data Fig. 9a). The impact of CH on HM mortality was more prominent when it was present in combination with abnormal blood counts, particularly cytopenia. A significantly higher HM mortality associated with CH was observed in subjects with abnormality in blood counts than in those without (Fig. 4e), depending on the number of CH-related lesions and on the severity of cytopenia; as large as 3.4% 10-year HM mortality was observed for those with multi-lineage cytopenia and multiple CH-related SNVs/indels and CNAs, compared with 0.46% for those with normal blood count lacking CH-related lesions.

The presence of both SNVs/indels and CNAs was associated with a significantly increased HM mortality compared with that of SNVs/indels (HR=2.84, 95%CI:2.14-3.78) or CNA (HR=2.64, 95%CI:1.94-3.60) alone (Fig. 4f). It was observed even when subjects were stratified according to the number of SNVs/indels (Extended Data Fig. 8c-e). However, the combined effect seems to be explained in large part by an increased total number of alterations, rather than the type of lesions co-occurred, i.e., SNVs/indels vs. CNAs. In fact, the HM mortality significantly correlated with the total number of CH-related lesions and the co-occurrence of both lesions did not significantly affect the mortality of individuals having the same number of lesions (Extended Data Fig. 8f-h). Many of SNVs/indels conferring a higher HM mortality, including those affecting *U2AF1, SRSR2, TP53*, and *JAK2*, tended to have a higher total number of CH-related lesions, compared with other SNVs/indels (Extended Data Fig. 2m). Nevertheless, the effect on HM mortality was not uniform across different combinations of SNVs/indels and CNAs, regardless of the total number of lesions. In particular, those involving the same gene/locus were associated with a higher HM mortality, compared with other combinations of SNVs/indels and CNAs (Extended Data Fig. 6c). The increase mortality was largely explained by those affecting *TP53*. However, even excluding *TP53*-involving SNVs/indels and CNAs, the combinations of lesions affecting the same locus showed a higher HM mortality than other SNVs/indels and CNAs combinations. Of interest, *TP53*-involving SNVs/indels also exhibited significant associations with del(5q) and multiple (≥3) CNAs mimicking a complex karyotype (Fig. 2a, Extended Data Fig. 6f), which together with 17pLOH, are among the most common lesions associated with *TP53* alterations in a variety of myeloid neoplasms with a very poor prognosis, particularly in MDS^25,33,41^. In agreement with this, these combinations involving *TP53* alterations were significantly associated with a higher mortality from MDS, compared with *TP53*-involving SNVs/indels alone (Extended Data Fig. 6g-h).

An almost identical risk estimation for HM was obtained in a case-control setting including all 672 cases who developed HM (Extended Data Fig. 1a, 9a). A small number of cases in which the onset of HM was recorded due to incomplete follow-up and exclusion of MDS and myeloproliferative neoplasms from the follow-up prevented powered analyses of the effect of CH on cumulative incidence of HM, although a similar trend of the effect of CH was observed with regard to the risk of HM that were seen in the analysis using mortality as an endpoint (Extended Data Fig. 9b-f).

### Effect of SNVs/indels and CNAs on cardiovascular mortality

Finally, we investigated the combined effect of SNVs/indels and CNAs on cardiovascular mortality in the cohort of 10,623 individuals using multivariate models to take into account known risk factors other than CH: age, gender, body-mass index, comorbidities (diabetes mellitus, hypertension, and dyslipidemia), history of smoking/drinking, and versions of SNP array. In accordance with the previous reports^13^, the presence of CH-related SNVs/indels with large clone size (VAFs ≥ 5%) were associated with an elevated cardiovascular and all-cause mortality (HR=1.36, 95%CI:1.09-1.71 for cardiovascular mortality; HR=1.41, 95%CI:1.24-1.60 for all-cause mortality) (Fig. 6a, Extended Data Fig. 10a). In support of this, we observed significant association of SNVs/indels with hypertension (Fig. 6b), which was independent of known risk factors for hypertension, including older age, a higher BMI, and diabetes. By contrast, regardless of their clone size, CNAs alone did not seem to affect cardiovascular or all-cause mortality (Fig. 6c, Extended Data Fig. 10b). However, CNAs in combination with SNVs/indels with ≥5% VAFs were significantly associated with elevated cardiovascular mortality and all-cause mortality, compared with CNAs alone, SNVs/indels alone and either SNVs/indels or CNAs (Fig. 6d and Extended Data Fig. 10c), although there was no significant difference in cardiovascular mortality or overall survival depending on whether or not they involved the same locus (Extended Data Fig. 6d,e). In multivariate analysis, the combined effect of both lesions was independent of the number of cooccurring SNVs/indels (HR=1.77, *P*=0.012) (Extended Data Fig. 10d-f). Given no impact of CNAs alone, the combined effect on cardiovascular and all-cause mortality does not seem to be explained by an increased total number of CH-related lesions. In fact, the total number of CH-related lesions did not correlate with cardiovascular and all-cause mortality, except for a significantly higher mortality for ≥3 CH-related lesions (Extended Data Fig. 10g), likely involving both SNVs/indels and CNAs. Collectively, these observations suggested that the presence of both SNVs/indels and CNAs increased the cardiovascular and all-cause mortality, compared with either of both lesions.

## Discussion

Combining targeted deep sequencing of major CH-relate genes and SNP array-based copy number analysis of blood-derive DNA from >10,000 individuals aged ≥60 years, we have delineated a comprehensive registry of CH in a general population of elderly individuals in terms of both SNV/indel and CNA. A case-cohort study design enabled an accurate estimation of CH-associated cumulative HM mortality in a large general cohort of elderly individuals (>43,000) including >400 cases who developed HM, substantially saving the cost and effort of sequencing, where only ∼8300 (∼18%) individuals/subcohort were fully genotyped. It should be noted that with a much larger number of cases with HM mortality (n=401) compared with previous cohort studies (16 and 37 cases/cohort)^12,13^, the estimation of HM mortality in individuals with CH-related SNV/indels was substantially more accurate with a much smaller confidential interval for both myeloid and lymphoid malignancies, where the mortality attributable to CH was mostly explained by myeloid malignancies regardless of type of CH-related lesions. Estimation of odds ratios for CH(+) vs. CH(−) cases were even more accurate with a total of 672 HM events in a case-control study setting.

Including both types of lesions, CH was found in as many as 40% of a general population of ≥60 years of age, of which 11% had ≥10% clone size. As a whole, SNVs/indels and CNAs co-occurred more frequently than expected only by chance. In particular, as repeatedly highlighted in myeloid neoplasms^29,33,42^, SNVs/indels in *DNTM3A, TET2, JAK2*, and *TP53*, significantly co-occurred with LOH at each locus in CH, suggesting the role of biallelic alterations of these genes even in an early stage during leukemogenic evolution. Co-occurrence of *TET2*-involving SNVs/indels and deletions involving the *TCRA* locus that are suggestive of evolution of *TET2*-mutated T-cell clones is also of interest. However, even excluding the subjects having these combinations affecting the same gene, SNVs/indels and CNAs significantly co-occurred (*P*=0.0042). Given that most of the CNAs in CH are recurrently seen in myeloid neoplasms, this suggests the presence of functional interactions between CH-related SNVs/indels and CNAs for positive selection, although we cannot exclude a possibility that CNAs might just represent chromosomal instability induced by one or more CH-related SNVs/indels.

Compared with those having SNVs/indels or CNAs alone, CH(+) individuals with both lesions showed a higher clone size, more abnormal blood counts, and a higher mortality from HM, particularly of myeloid lineages. The combined effect of SNVs/indels and CNAs^40^, is typically exemplified by biallelic alterations in *DNTM3A, TET2, JAK2*, and *TP53*, caused by LOH affecting the mutated locus. However, the effect of combined SNVs/indels and CNAs is largely explained by an increased total number of CH-related lesions. Given that the size of CH clones correlated with the number of CH-related lesions, the increasing number of mutations is thought to promote expansion of clones, contributing to an earlier onset and progression of HM. This underscores the importance of measuring both lesions for accurate estimation of HM mortality, which is expected to increase the number of CH-related lesions evaluated only for SNVs/indels and CNAs alone by 0.25 and 0.36 on average, revising 10-year expected HM mortality by 0.14% and 0.19%, respectively. The combined effect of both SNVs/indels and CNAs was also observed for cardiovascular and all-cause mortality. Of interest, the effect was seen despite that CNAs alone did not affect the mortality. Because the effect of SNVs/indels on cardiovascular mortality depended on their VAFs, which increased with the presence of CNAs, the combined effect seems to be mediated in part by an increased size of clones having SNVs/indels, although CNA still remained significant after the effects of clone size was adjusted.

Potential caveats in the current study include a limited number of CH-related genes analyzed (n=23), a compromised sensitivity of detecting focal CNAs, and the study population exclusively including individuals over 60 years of age. However, these 23 genes, which are estimated to capture ∼90% of CH-related SNVs/indels^12,13^, were analyzed using deep sequencing to sensitively detect lesions in very small fractions (∼1%), which would not have been possible with a more unbiased sequencing with a larger target size. In addition, CH and related HM and CVD are highly enriched in and mostly confined to this age group, respectively. Thus, the limited number of genes and age group might not necessarily be the limitations, but rather contributed to efficient analyses of comprehensive analysis of CH-related alterations in a large number of cases to investigate their effects on clinical outcomes at an acceptable cost. However, clearly more comprehensive studies with unbiased sequencing and improved copy number detection including all age groups should be warranted to elucidate the full spectrum of CH-related alterations in future studies.

## Acknowledgement

This work was supported by the Japan Agency for Medical Research and Development (AMED) (JP15cm0106056h0005, JP19cm0106501h0004, JP16ck0106073h0003, JP19ck0106250h0003 to S.O.; JP17km0405110h0005 and JP19ck0106470h0001 to H.M.; JP19ck0106353h0003 to Y.N.) and the Core Research for Evolutional Science and Technology (CREST) (JP19gm1110011 to S.O.); the Ministry of Education, Culture, Sports, Science and Technology of Japan; the High Performance Computing Infrastructure System Research Project (hp160219, hp170227, hp180198 and hp190158 to S.O. and S.M.) (this research used computational resources of the K computer provided by the RIKEN Advanced Institute for Computational Science through the HPCI System Research project); the Japan Society for the Promotion of Science (JSPS); Scientific Research on Innovative Areas (JP15H05909 to S.O. and S.M.; JP15H05912 to S.M.) and KAKENHI (JP26221308 and JP19H05656 to S.O.; JP16H05338 and JP19H01053 to H.M.; JP15H05707 to S.M.); the Takeda

Science Foundation (S.O., H.M. and T.Y.). S.O. is a recipient of the JSPS Core-to-Core Program A: Advanced Research Networks. DNA samples and subjects’ clinical data were provided by Biobank Japan, the Institute of Medical Science, the University of Tokyo. The super-computing resource was provided by Human Genome Center, the Institute of Medical Science, the University of Tokyo.

## Author contributions

R.S., H.M., and S.O. designed the study. K.M., Y.K., T.M., and Y.M. provided DNA samples and clinical data. Y.K. and S.M. provided bone marrow samples. C.T., and Y.K. performed copy-number analysis. Y.M. and M.K. performed sequencing. M.M.N. performed cell sorting and single-cell analysis. R.S., M.M.N., Y.O., T.Y., Y.S, K.C., H.T., A.N., S.I., and S.M. performed bioinformatics analysis. R.S., Y.N., M.M.N., Y.O., T.Y., H.M., and S.O. prepared the manuscript. All authors participated in discussions and interpretation of the data and results.

## Online methods

### Sample ascertainment

All subjects in this study were derived from BioBank Japan (BBJ) project, a multi-hospital-based-registry^23^. BBJ project enrolled approximately 200,000 individuals with at least one of 47 target diseases between fiscal years 2003 and 2007. From 179,417 participants of BBJ project in which SNP array analysis of peripheral blood-derived DNA had been performed, we enrolled a total of 11,234 subjects. Among these, 10,623 were randomly selected from 60,787 cases who were aged ≥60 years at the time of sample collection and were confirmed not to have solid cancers as of March 2013. Out of the randomly selected 10,623 cases, 61 were recorded to develop or die from HM. The remaining 611 subjects, all of whom were recorded to have HM events, were additionally enrolled to maximize the statistical power in survival analysis. In total, we enrolled 672 subjects with any HM events, 138 and 589 of which were recorded to develop and die from HM, respectively. Subjects’ demographic summary was presented in Supplementary Table 1. The numbers of subjects with individual targeted diseases were listed in Supplementary Table 2. The protocols for this study were approved by the ethics committees at all the involved institutions; written informed consent had been obtained from all participants.

### Multiplex PCR-based targeted sequencing

To detect CH-associated driver mutations, we performed multiplex PCR-based targeted sequencing, as previously described.^24^ Primers were designed to cover coding regions of 23 driver genes commonly mutated in clonal hematopoiesis or myeloid neoplasms: *ASXL1, CBL, CEBPA, DDX41, DNMT3A, ETV6, EZH2, GATA2, GNAS, GNB1, IDH1, IDH2, JAK2, KRAS, MYD88, NRAS, PPM1D, RUNX1, SF3B1, SRSF2, TET2, TP53*, and *U2AF1*. PCR product sizes were designed to be 180-300 bp to cover the amplicon by the sequencing reads. We added CGCTCTTCCGATCTCTG to the 5’ end of the forward primers and CGCTCTTCCGATCTGAC to the 5’ end of the reverse primers to perform second PCR.^43,44^ We performed multiplex PCR using different primer pools to cover all coding regions of the 23 genes. Then we performed second PCR with primer sequences 5’-AATGATACGGCGACCACCGAGATCTACACxxxxxxxxACACTCTTTCCCTACACGACGCTCTTCCGATCTCTG-3’ and 5’-CAAGCAGAAGACGGCATACGAGATxxxxxxxxGTGACTGGAGTTCAGACGTGTGCTCTTCCGATCTGAC-3’, where xxxxxxxx represents 8-bp barcodes. All second PCR products were pooled for one sequencing run. After each library was purified using Agencourt AMPure XP (Beckman Coulter), we obtained 2×150-bp paired-end reads with dual 8-bp barcode sequences on a HiSeq2500 instrument.

### Calling CH-related SNVs/indels

Sequencing reads were aligned to the human genome reference (hg19) using Burrows-Wheeler Aligner, version 0.7.8, with default parameter settings. Mutation calling was performed through our established pipeline, as previously reported^33,45-47^, using the following parameters.

First, we adopted variants fulfilling the following criteria:

i. Number of variant reads ≥ 10 (≥5 for TCGA dataset) †
ii. Variant allele frequency (VAF) ≥ 0.5%†
iii. Non-synonymous variants within coding-sequence or splice-site variants

(† For calculation of read counts and VAFs, we only counted base calls fulfilling Mapping Quality score ≥ 40, and Base Quality score ≥ 20.)

To further exclude false positive calls due to sequencing artifacts, we modeled site-specific error rates as beta-binominal distribution. Parameters for beta-binomial distribution were determined by maximum likelihood method^48^, based on the read counts in all samples. Mutation calls whose VAFs were significantly deviated from background-error distribution (*P*beta-binomial ≤ 10^−6^) were regarded as true mutations.

Additionally, variants always appeared within similar ranges of VAFs (especially <1%, or >40%) were likely to be sequencing artefacts or germline polymorphisms, rather than true somatic mutations. Based on this assumption, we excluded candidates fulfilling both of the following criteria from the remaining candidates:

i. Candidates observed in ≥5 samples
ii. Mean VAF <1%, or >40%, or coefficient of variation of VAFs < 0.5.

The candidates fulfilling the quality filter noted above were included in the subsequent analyses if they fulfil one of the following criteria for driver mutations^33^:

i. Candidates resulting in amino-acid substitutions which were registered in the Catalogue of Somatic Mutations in Cancer (COSMIC) v91 databases for ≥ 5 counts
ii. Candidates which fulfill the Criteria 1 and at least one of the Criteria 2

Criteria 1

Candidates which were not registered in public databases, including dbSNP138, the 1000 genomes project as of 2014 Oct, Human Genome Variation Database, and The Exome Aggregation Consortium (ExAC).

Criteria 2

i. Candidates located on the non-repeat region with VAFs ≥4% <40% or ≥60% <96%
ii. Nonsense, frameshift, or splice-site candidates
iii. Candidates which were computationally predicted to have negative consequences by SIFT (score < 0.05), PolyPhen-2 (damaging), and MutationAssessor (high or medium)

Finally, the resulting set of driver mutations were manually reviewed in Integrated Genome Viewer (http://software.broadinstitute.org/software/igv/).

### *In silico* simulation of mutation calling

To benchmark the performance in detection of low-VAF mutations, we performed *in silico* simulation. Mixing 2 bam files with variable proportions, we diluted 750 heterozygous SNPs and artificially created low-VAF mutations (ranging from 0.5% to 5%). Each diluted SNPs were classified into 6 bins according to sequencing depths (x100-x300, x300-x500, x500-x750, x750-x1000, x1000-1500, and x1500-), and sensitivities were calculated separately for the 6 bins. We calculated sensitivity as a fraction of detected variants within all simulated variants:

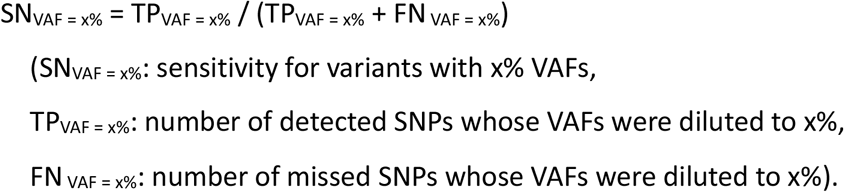

Together with sensitivity, we calculated specificity by sampling genomic positions without known SNPs (n = 5000/simulation). We counted mutation calls on these positions as false positives, and calculated the specificity as follows:

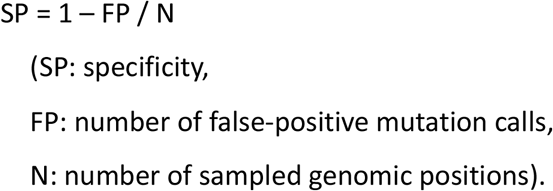

To draw receiver operator characteristic (ROC) curves, we calculated sensitivities and specificities for 9 different cutoffs on beta-binomial *P* values (10^−2^, 10^−3^, 10^−4^, 10^−5^, 10^−6^, 10^−7^, 10^−8^, 10^−9^, and 10^−10^).

### Copy-number analysis

Our analysis pertaining CNAs are based on the result in previous publication^21^, in which blood derived DNA samples from the 11,234 subjects were examined by either of three different versions of microarrays: Illumina Infinium OmniExpress (n=708), Infinium OmniExpressExome v.1.0 (n=3,152), or v.1.2 (n=7,374). For detection of CNAs, we analyzed allele-specific hybridization intensities for the polymorphisms examined by all versions of arrays (n = 515,355). Haplotype phasing and calculation of log R ratio (LRR) and B-allele frequency (BAF) were performed as previously described^21^. Based on long-range haplotype information and LRR/BAF values, we detected allelic imbalances and classified them into duplications, deletions, and UPDs, with false discovery rate around 5%.^10,21^ Because the power to detect allelic imbalances exceeded the power to distinguish UPD from copy-number gain or loss, CNAs were designated as “unclassifiable” when we could not assign them into specific types of CNAs. In the analyses where the exact discrimination between UPD, duplication, or deletion (e.g., lesion-specific analysis in Fig 2a, 3) was relevant, we excluded unclassifiable CNAs from the analysis. Although we cannot calculate precise cell fractions for unclassifiable CNAs, their cell fractions are basically expected to be quite small. Therefore, when we classified CNAs by their cell fractions (e.g., Fig. 4d, 5b, and 6c), unclassifiable CNAs were regarded to be smaller than the thresholds. When we analyzed CNAs in terms of their cell fractions (e.g., Fig. 1d,e, 5a), unclassifiable CNAs were excluded. Otherwise, we did include those unclassifiable CNAs in the analysis (e.g., Fig 1a-c, 2b-e, 3c-d, 4, 6b,d). Based on the detected CNAs, we determined chromosomal regions significantly affected with CNAs by PART (parametric aberration recurrence test)^49^ (https://www.hgc.jp/~aniida/PART/manual.html).

### Definition of abnormalities in blood counts

Subjects fulfilling at least one of the following criteria were considered to have abnormalities in blood counts.

i. White blood cells (/µL): ≥10000, or <3000
ii. Hemoglobin (g/dL): ≥16.5 (male), ≥16 (female), or <10
iii. Hematocrit (%): ≥50
iv. Platelet (10000/µL): ≥50, or <10

These cutoffs on blood counts were adopted from diagnostic criteria for MDS or myeloproliferative neoplasms^50^. Out of subjects with available counts for all of WBC, hemoglobin, hematocrit, and platelet (n = 8,345), 7,031 subjects (84.3%) had normal blood cell counts.

### Analysis of lineage-sorted samples

Bone marrow Frozen bone marrow was thawed in Dulbecco’s Modified Eagle Medium (Sigma-Aldrich) containing 10% of foetal bovine serum (FBS, biosera) and 1% of Penicillin-Streptomycin solution (ThermoFisher). After the cell pellets were washed with PBS containing 2% FBS, the cells were stained with an antibody mix for 20 min, followed by washing with PBS containing 2% FBS and filtered with a 5 mL Round Bottom Polystyrene Test Tube with Cell Strainer Snap Cap (ThermoFisher). We mixed 500 µL of the filtered cell suspension in PBS containing 2% FBS was mixed with 5 µL of Propidium Iodide Staining Solution (BD Bioscience), which was then sorted with the FACSAria III cell sorter (BD Bioscience). The antibodies used in flow cytometry are listed in Supplementary Table 4. For digital droplet PCR (ddPCR) and amplicon-sequencing, we sorted myeloid, erythroid, T cell, and B cell fractions and gDNA was extracted from sorted cells. To detect allelic imbalances in the region of del(13q), amplicon sequencing was performed with custom primers targeting heterozygous SNPs within the deleted region (ThermoFisher, Supplementary Table 5). To detect the A1153V substitution in *TET2*, ddPCR was performed as described below. For single-cell analysis, CD34^+^ cells were sorted. Cells were re-suspended in StemSpan Serum-Free Expansion Medium (STEMCELL Technologies) at 400–1,600 cells/µL, which was then applied into Fluidigm C1 platform for combined single-cell gene expression analysis and SNV detection. Detailed methods for single cell analysis are in preparation for publication (shared upon request, Masahiro M Nakagawa, Ryosaku Inagaki, et al.).

### ddPCR

For ddPCR, predesigned probes were purchased from BioRad. We mixed 50 ng of gDNA with enzymes (ddPCR Supermix for Probes (no dUTP), BioRad) and the probe mix, followed by droplet generation and PCR amplification according to the manufacturer’s protocol. Annealing temperatures was set at 55°C. We measured amplified droplets using the QX200 system and QuantaSoft (version 1.7, BioRad). Catalogue numbers of probe mix are shown in Supplemental Table 6.

## Statistical analysis

All the statistical analyses were performed using the R statistical platform (https://www.r-project.org/) v.3.6.1. All statistical tests were two-sided. Benjamini–Hochberg multiple testing correction was applied when appropriate.

### Age-stratified permutation test for cooccurrences of CH-related alterations

We tested the significance of cooccurrences between SNVs/indels and CNAs under the stratification by subjects’ age, because age-dependent frequencies of both CH-related alterations can confound their cooccurrences. First, we stratified subjects into 41 bins according to their age (60, 61, …, 100 years old) and calculated frequencies of SNVs/indels, CNAs, and their cooccurrences within each bin. In single iteration of permutation, we randomized the status of SNVs/indels and CNAs in all subjects while retaining their frequencies in each age bin. Then, the number of cooccurrences were summed up across all age bins. By repeating this process, we obtained null random distribution of the number of subjects with cooccurring SNVs/indels and CNAs. Comparing the null distribution and the actual number of cooccurrences, we obtained *P* value for significance of cooccurrences between SNVs/indels and CNAs. Significant cooccurrences of multiple CH-related alterations was also demonstrated in a similar way, in which we counted the total number of CH-related alterations within each age bin. In single iteration, these alterations were randomly re-assigned to the subjects retaining the total number of alterations in each bin. Then, the number of subjects to whom multiple alterations were assigned was counted across all bins. *P* value was calculated by comparing the actual number of cases with ≥2 alterations and null distribution generated by repeating the process above.

### Simulation test for cell-level coexistence of SNVs/indels and CNAs involving the same genes

Regarding the combinations of SNVs/indels and UPDs involving the same genes (*DNMT3A, TET2, TP53*, and *JAK2*), we observed higher VAFs of SNVs/indels than cell fractions of CNAs in 49 of the 51 cases, which suggested they were likely to be acquired in the same cells and resulted in biallelic alterations (Supplementary Figure 6a,b). To examine how many of the 49 cases should be explained by cell-level coexistence of SNVs/indels and UPDs, we performed random simulation on their clone sizes putting a null hypothesis, *H*_0_(x): SNVs/indels and UPDs were independently acquired in at least x cases (x=3,4,…,51). *P* value for *H*_0_(x) was calculated assuming VAFs of SNVs/indels and cell fractions of UPDs follows independent distributions (Supplementary Figure 6c-e). We searched for the maximum x with which *P* value for *H*_0_(x) was below 0.05 to obtain minimum estimate of the number of cases in which cell-level coexistence of SNVs/indels and UPDs was expected (Supplementary Figure 6f).

### Risk factors for CH

To extract risk factors for CH, we examined correlations between genetic alterations in CH and baseline characteristics of subjects (age, sex, history of smoking and drinking). Information regarding the history of smoking and drinking were based on self-report questionnaires at DNA sampling. First, we performed univariate logistic regressions for presence of genetic alterations. Based on factors significantly correlated with genetic alterations (*q* < 0.1), we then performed multivariate logistic regressions to extract independent risk factors (*P* < 0.05).

### Effect of CH on blood cell counts

To elucidate effects of genetic alterations on blood cell counts, we examined correlations between genetic alterations and blood cell counts. After Cox-Box transformation of blood counts, linear regressions were performed. To correct for confounding effects, all regressions were perfumed in multivariate models including age, gender, and versions of SNP array as covariates, in comparison with subjects without detectable CH.

### Prediction models for hypertension

To elucidate the relationships between CH and hypertension, we performed multivariate logistic regression. Optimal sets of variables were selected by stepwise method from known risk factors and blood test values available for ≥70% of the subjects: presence of SNVs/indels and CNAs, age (+10 years), gender, BMI (+5), history of smoking and drinking (based on self-report questionnaires), white blood cell count, red blood cell count, hemoglobin, hematocrit, MCHC, platelet, aspartate aminotransferase (AST), alanine aminotransferase (ALT), lactate dehydrogenase (LDH), creatinine, blood urea nitrogen, total cholesterol, and glucose.

### Survival analysis

We evaluated the effects of CH-related mutations, CNAs, and their combinations on mortality from HM, all-cause mortality, and cardiovascular mortality. To define mortality from hematologic malignancies, we included diagnoses within ICD10 code groups C81–C96 (malignant neoplasms of lymphoid, hematopoietic and related tissue), D45 (polycythemia vera), D46 (MDS), D47 (other neoplasms of uncertain behavior of lymphoid, hematopoietic and related tissue), and D7581 (myelofibrosis). For CVD, we included I20-I25 (ischemic heart diseases), I48-49 (arrythmia), I50 (heart failure), I60-I67, I69 (cerebrovascular diseases), I70-I72 (aortic atherosclerosis, aortic aneurysm, aortic dissection), and I74 (peripheral artery diseases). In analysis on all-cause mortality, we performed Cox proportional hazards regression using the R package, “survival” (http://cran.r-project.org/web/packages/survival/index.html). In analysis of HM events (mortality or development) or mortality from CVD, we performed competing risk regression based on fine-gray model. In the analysis of events of HM (mortality and development), we applied a case-cohort design to maximize the statistical power as previously described^38^ (Extended Data Fig. 1b, 9b), including all subjects with HM events within the target cohort. Meanwhile, cardiovascular mortality and overall survival were analyzed in a cohort of the randomly selected 10,623 subjects. To correct for confounding effects, we included subjects’ age, gender and version of SNP array in the multivariate models for events of HM, while age, gender, BMI, presence of diabetes mellitus, hyperlipidemia, and hypertension, history of tobacco smoking and alcohol drinking, and version of SNP array were included in the models for all-cause and cardiovascular mortalities.

## Data availability

A table of detected somatic mutations is available at https://github.com/RSaikiRSaiki/CH_2020. Clinical data and a list of CNAs can be provided by the BBJ project upon request (https://biobankjp.org/english/index.html). All other data will be made available upon request to the corresponding author.

## Code availability

All computational codes are available upon request to the corresponding author.

**Extended Data Fig. 1.**
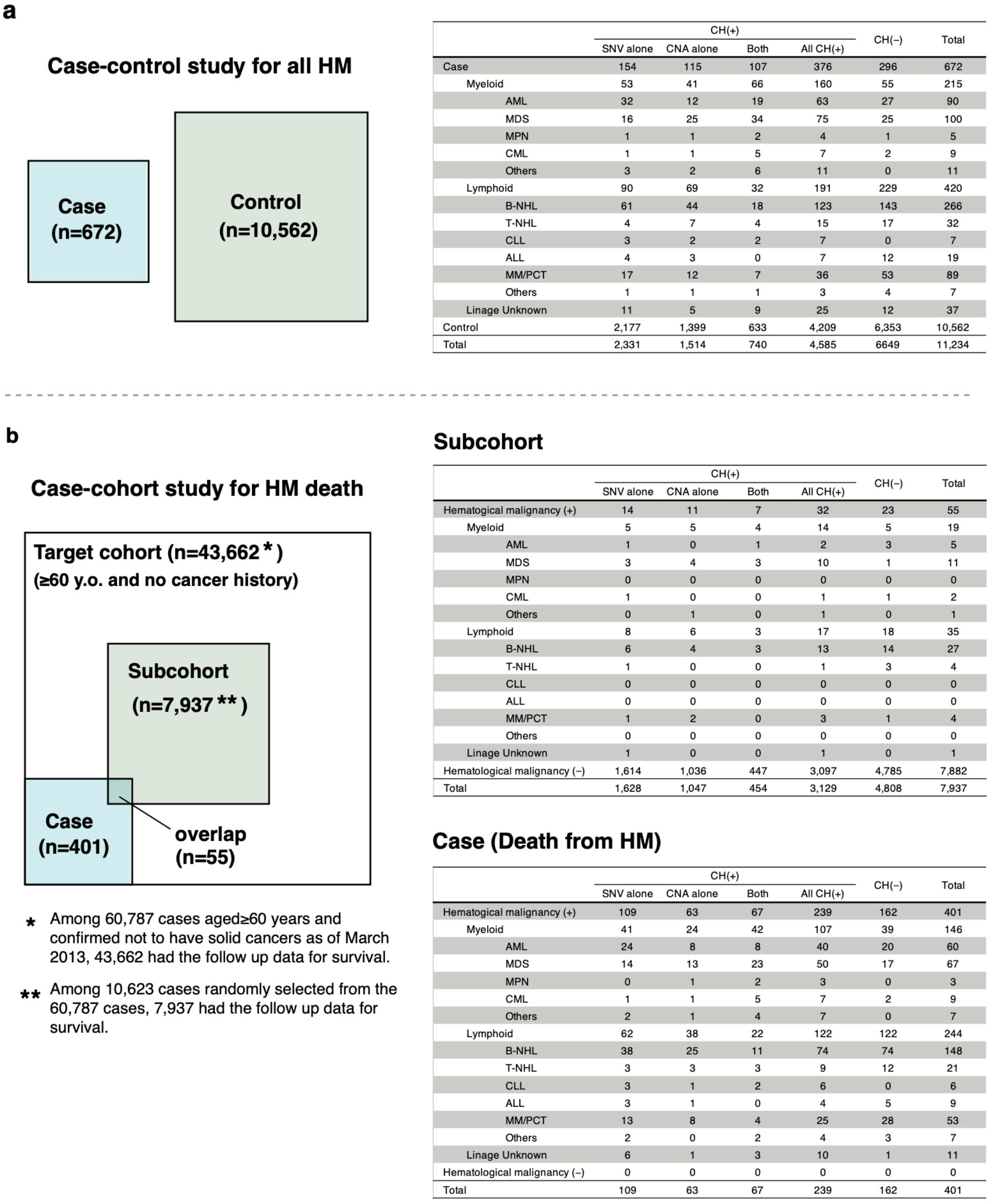
Design of case-control and case-cohort study. a, Design of case-control study (Left). Diagnosis of hematological malignancies (HM) in subjects with or without CH enrolled in the case-control study (Right). b, Design of case-cohort study for death from HM (Left). Diagnosis of HM in subjects with or without CH enrolled in the case-cohort study (Right). AML, acute myeloid leukemia; MDS, myelodysplastic syndromes; MPN, myeloproliferative neoplasms; CML, chronic myeloid luekemia; B-NHL, B-cell non-Hodgkin lymphoma; T-NHL, T-cell non-Hodgkin lymphoma; CLL, chronic lymphoid leukemia; ALL, acute lymphoblastic leukemia; **MM**, maltiple myeloma; PCT, plasma cell tumor.

**Extended Data Fig. 2.**
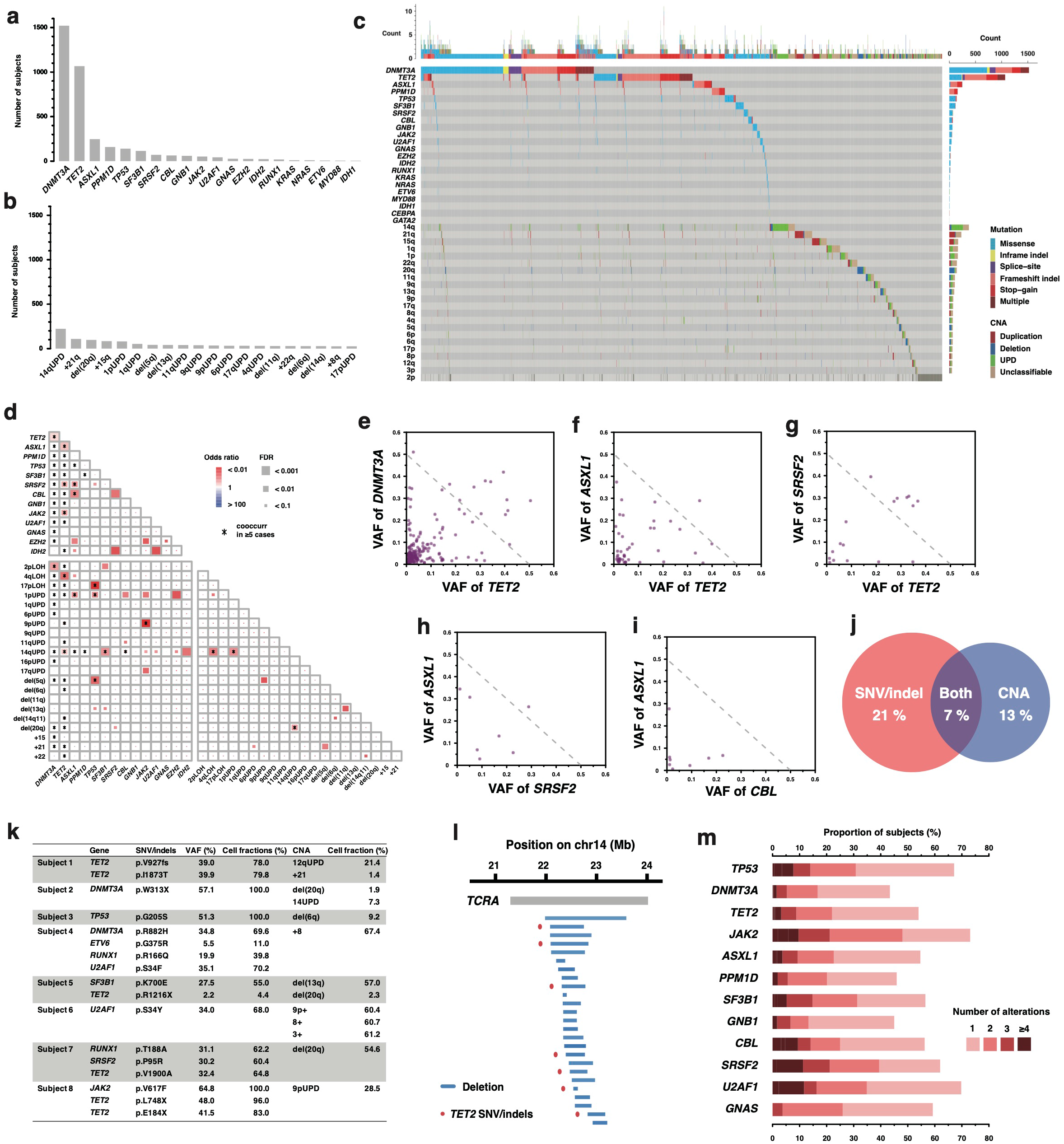
Landscape of genetic alterations in CH. a-b,The number of subjects with inidividual SNVs/indels (a) and CNAs (b). The vertical axis represents the number of subjects with indicated alterations. Unclassifiable CNAs are not included in (b). c, Landscape of SNVs/indels and CNAs in 11,234 subjects. Those without CH-related alterations are omitted. d, The correlations between individual genetic alterations. Combinations seen in 5 or more cases are indicated by asterisks. e-i, VAF of cooccurring SNV/indels in diagonal plot. Dots above the dashed line fufill “pegeonhall principle”. j, Venn diagram illustrating the overlap between subjects with SNV/indels and those with CNAs. Frequencies within all subjects in whom SNVs/indels and CNAs were examined (n=11,234) are indicated. k, Subjects in whom coocurring SNVs/indels and CNAs were suspected to coexist in the same cells on the basis of “pegeonhall principle.” I, A magnified illustration of microdeletions around *TCRA* locus (14q11.2). A gray bar represents gene body of *TCRA*. Blue horizontal bars represent microdeletions. Cooccurring *TET2* SNVs are indicated by red dots. Genomic coordinates in hg19 are indicated above. m, Proportions of subjects with different number of cooccurring alterations within those who harbor SNVs/indels in the indicated genes. The proportions of subjects with 1,2,3, and ≤4 CNAs are depicted by different colors.

**Extended Data Fig. 3.**
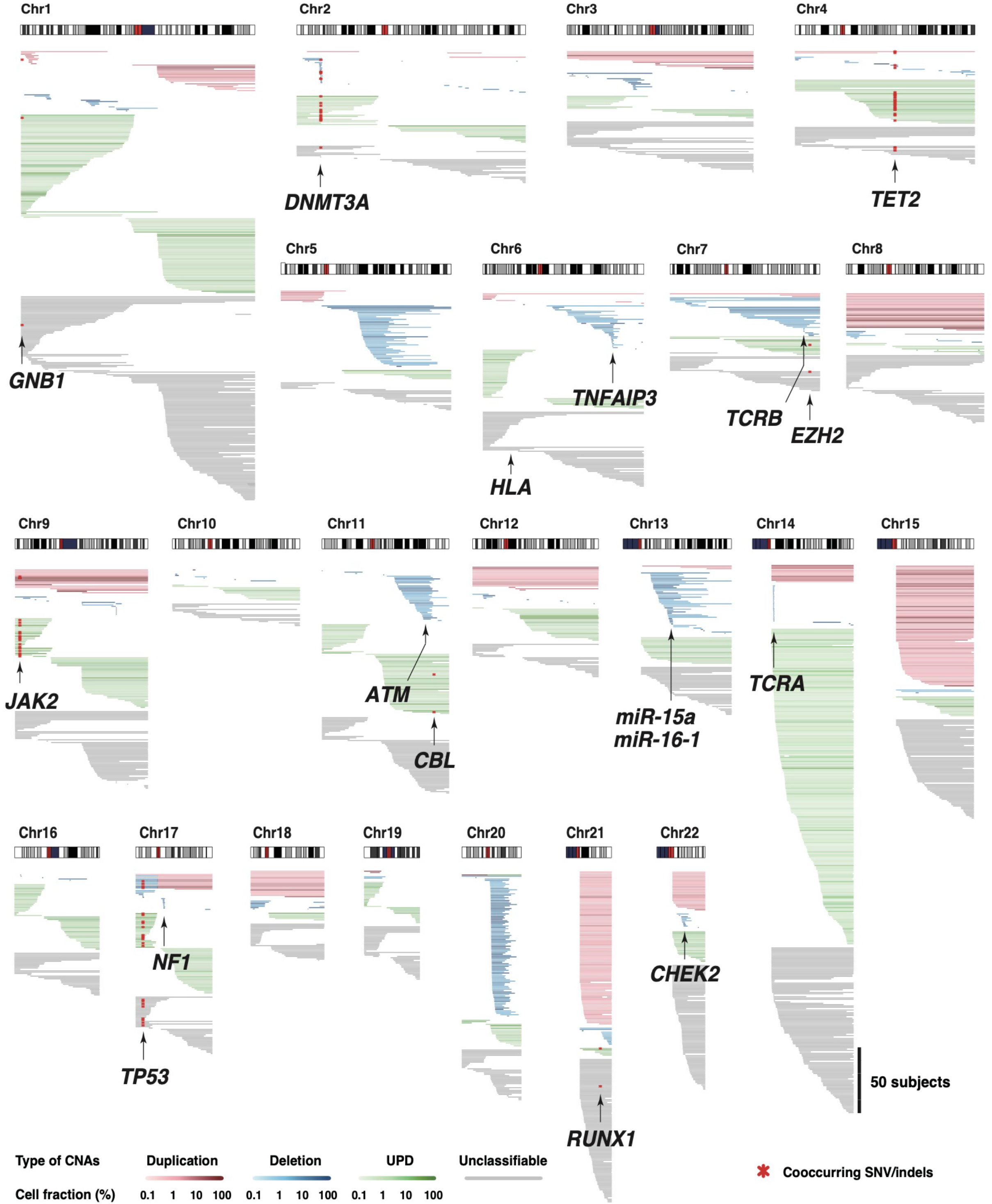
Distribution of CNAs in all chromosomes. Distributions of CNAs on all chromosomes are illustrated. Loci of known driver genes are indicated by arrows. Each horizontal bar represents one CNA. Cooccurring SNV/indels are indicated by red dots. Types of CNAs are depicted by different colors as indicated in the annotations.

**Extended Data Fig. 4.**
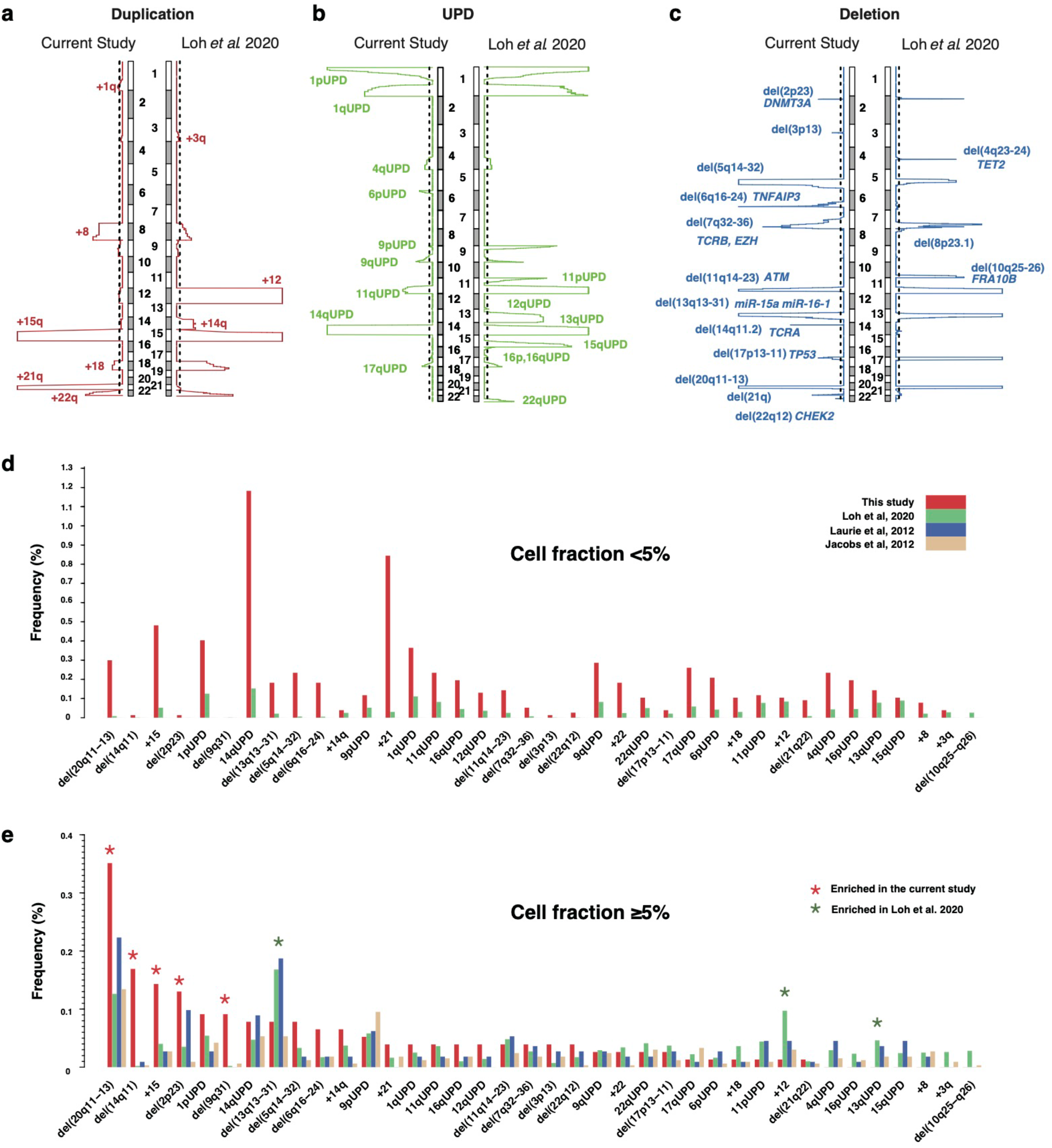
Chromosomal regions significantly affected by CNAs. a-c, Chromosomal regions significantly affected by duplications (a), UPDs (b), and deletions (c) in Japanese cohort (current study) and in British cohort”. Statistical significance for recurrence of CNAs were evaluated by PART”. Dashed lines indicate thresholds for statistical significance (q = 0.25). d-e, Comparison of frequencies of individual CNAs between the current and previous studies”. Comparisons were performed in those aged 60-75 years. In (d) or (e), CNAs in <5% or ≤5% cell fractions were taken into account, respectively. CNAs significantly enriched in either cohort were indicated by asterisks (q <0 .1) in (e).

**Extended Data Fig. 5.**
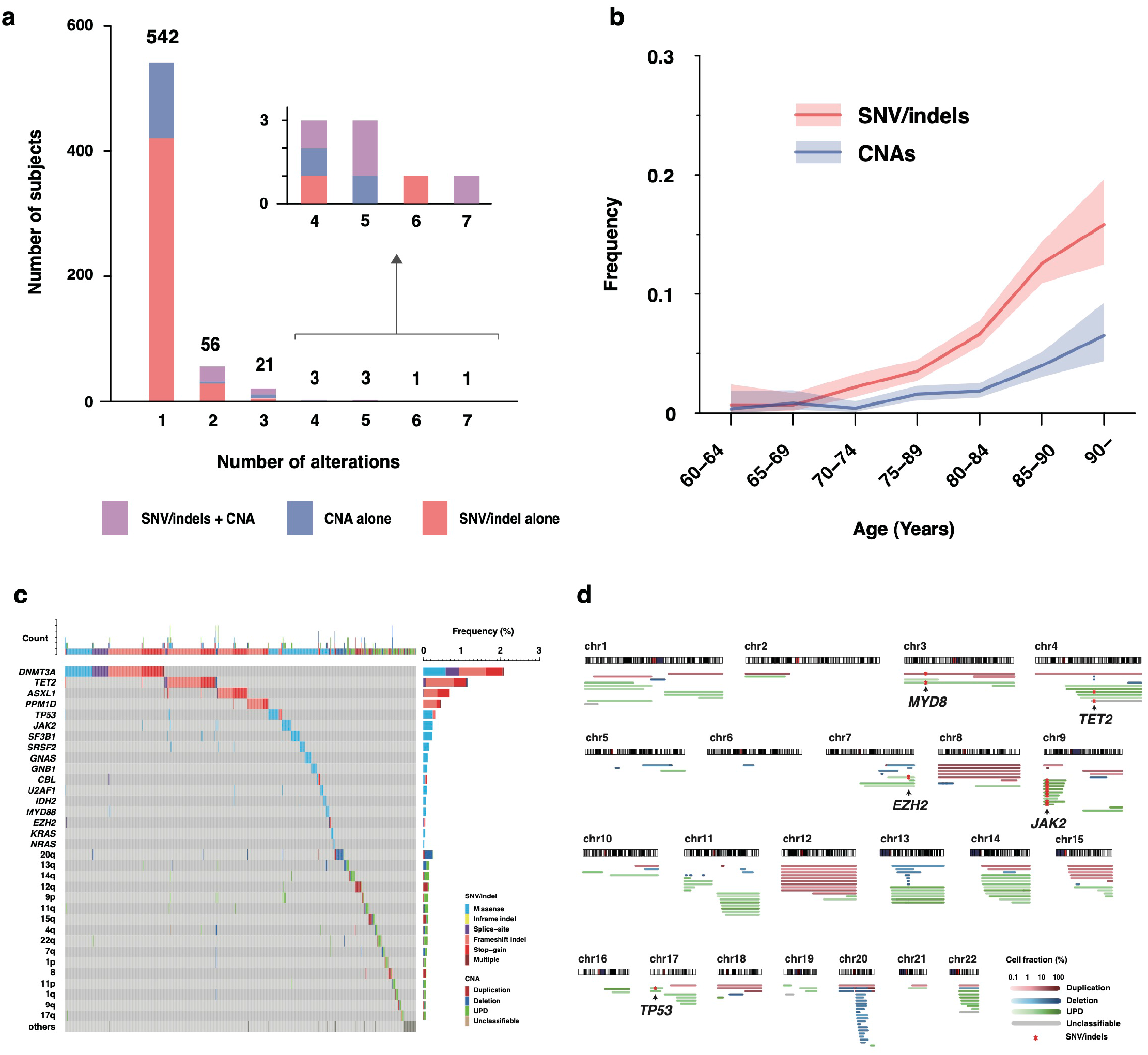
Analysis of SNVs/indels and CNAs in peripheral blood samples in TCGA cohort. a, Distribution of the number of genetic alterations in each subject. Subjects with SNVs/indels alone, with CNAs alone, or with both of them are illustrated by different colors. b, The prevalence of CH-related SNVs/indels and CNAs, according to age. Colored bands represent the 95% confidence intervals. c, The landscape of CH-related SNVs/indels and CNAs. Each row represents genetic alterations or affected chromosomal arms, and each column represents subjects. Subjects without any alterations are omitted. Types of SNVs/indels and CNAs are depicted by different colors. d, Distributions of CNAs on all chromosomes are illustrated. Loci of cooccurring SNVs/indels are indicated by arrows. Each horizontal bar represents one CNA. Cooccurring SNVs/indels are indicated by red asterisks. Types of CNAs are depicted by different colors.

**Extended Data Fig. 6.**
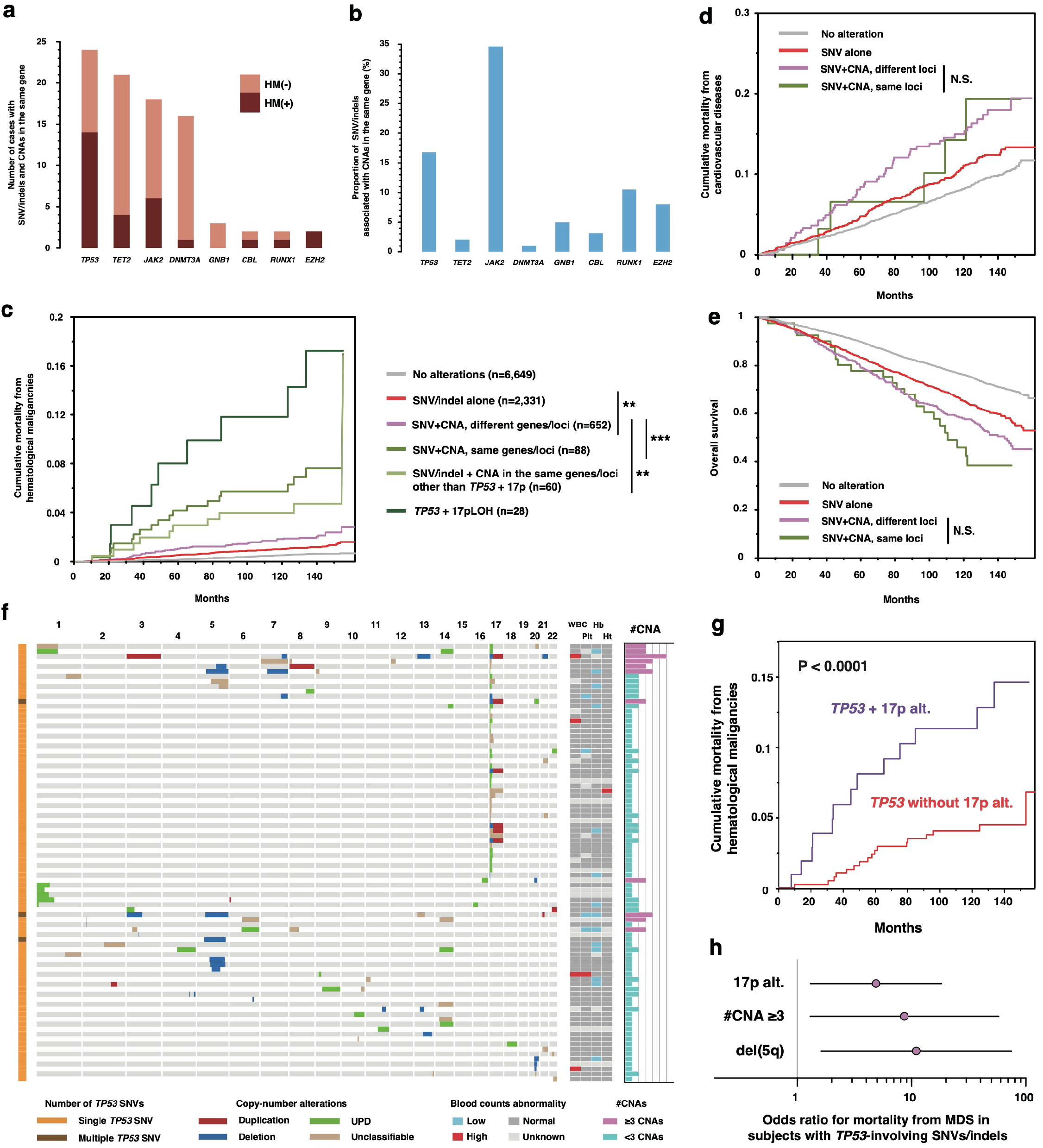
Interplay between SNVs/indels and CNAs. a, Number of subjects with SNVs/indels and CNAs involving the same genes/loci. b, Proportion of SNVs/indels associated with CNAs in the same genes/loci. c, Cumulative mortality from hematological malignancies. d, Cumulative mortality from cardiovascular diseases. e, Survival curves for overall survival. f, Profiles of CNAs in subjects with SNV/indels in *TP53*. Abnormaly high or low blood counts (WBC, Platelet, Hemoglobin, and Hematcrit) are indicated by red or blue, respectively. Numbers of coocurring CNAs are indicated on the right side (#CNA), where subjects with ≤3 CNAs were highlighted by purple. Subjects without any CNA were abbreviated. g, Mortality from hematological maligancnies in *TP53*-mutated cases with or without CNAs in 17p. h, Odds ratio for mortality from MDS calculated by multivariate logistic regression in subjects with *TP53*-involving SNVs/indels. We included unclassifiable CNAs involving 17p in 17p alterations (17p alt.) in panel (g-h) because they are most likey to be LOH (UPDs or deletions). *TP53*-involving SNVs/indels in panel (f-h) included those detected by ddPCR (Supplementary Fig. 3). N.S., not significant; **, *P* < 0.001; \*\*\**P* < 0.0001.

**Extended Data Fig. 7.**
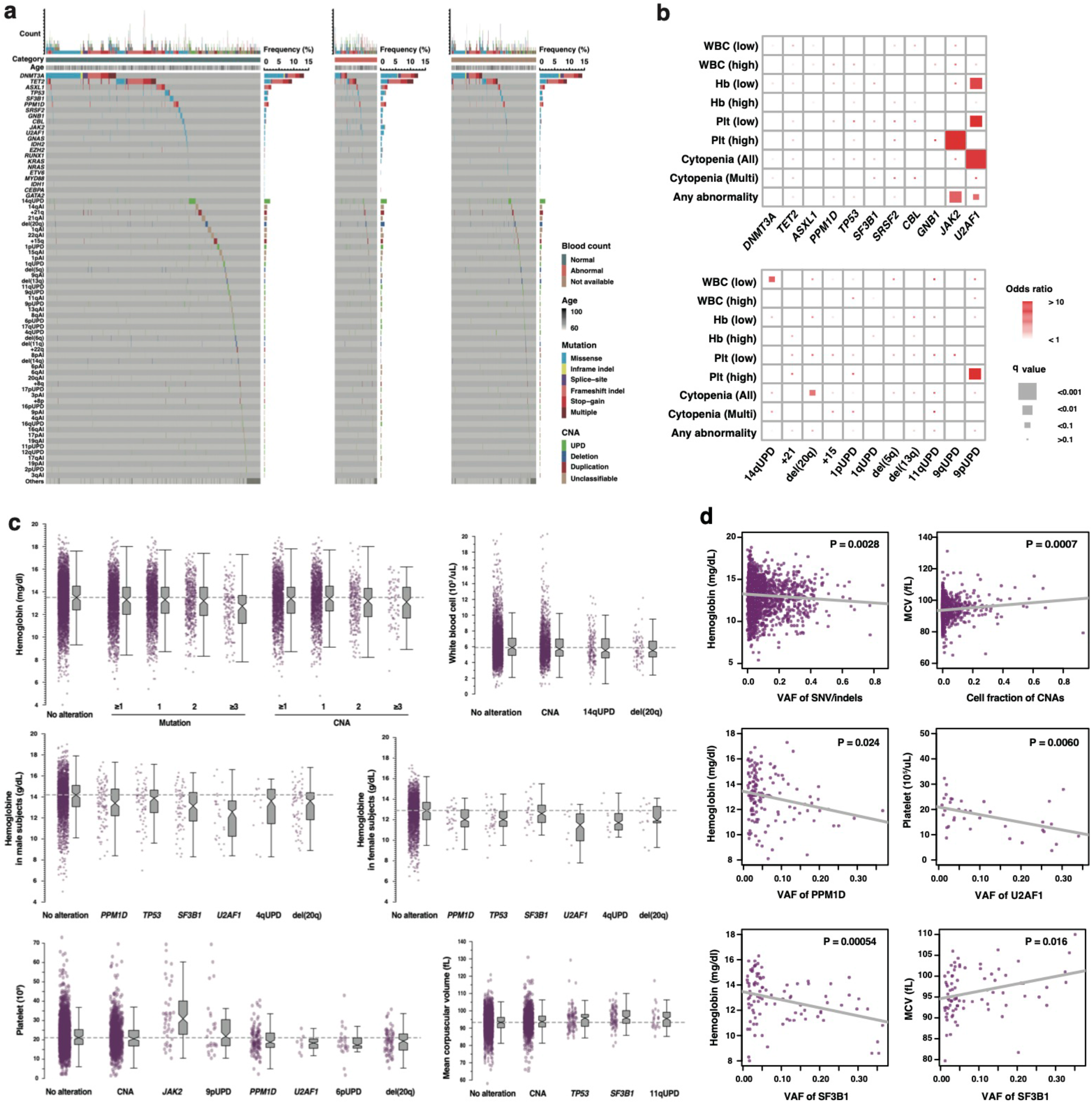
Genetic alterations In CH and abnormalities in blood counts. a, Landscape of SNVs/indels and CNAs in subjects without abnormalities in blood counts (left), in those with any abnormalities in blood counts (middle), and in those with no available blood counts (right). Each row represents a genetic alteration while each column represents a subject. Subjects without any alteration are omitted. Different types of mutations and CNAs are depicted by different colors. b, Enrichment of genetic alterations in subjects with abnormalities in blood counts. Sizes of rectangles indicate significance of enrichment. Colors of rectangles indicate odds ratios. The enrichment of alterations were examined by Fisher exact test. Cytopenia (All), subjects with cytopenia in at least one lineage; Cytopenia (Multi), subjects with cytopenia in 2≤ lineage. WBC, white blood cell; Hb, hemoglobin; Plt, platelet. c, Distribution of blood cel counts in subjects with different CH-related alterations. d, Relationships between blood cell counts and VAF of SNVs/indels.

**Extended Data Fig. 8.**
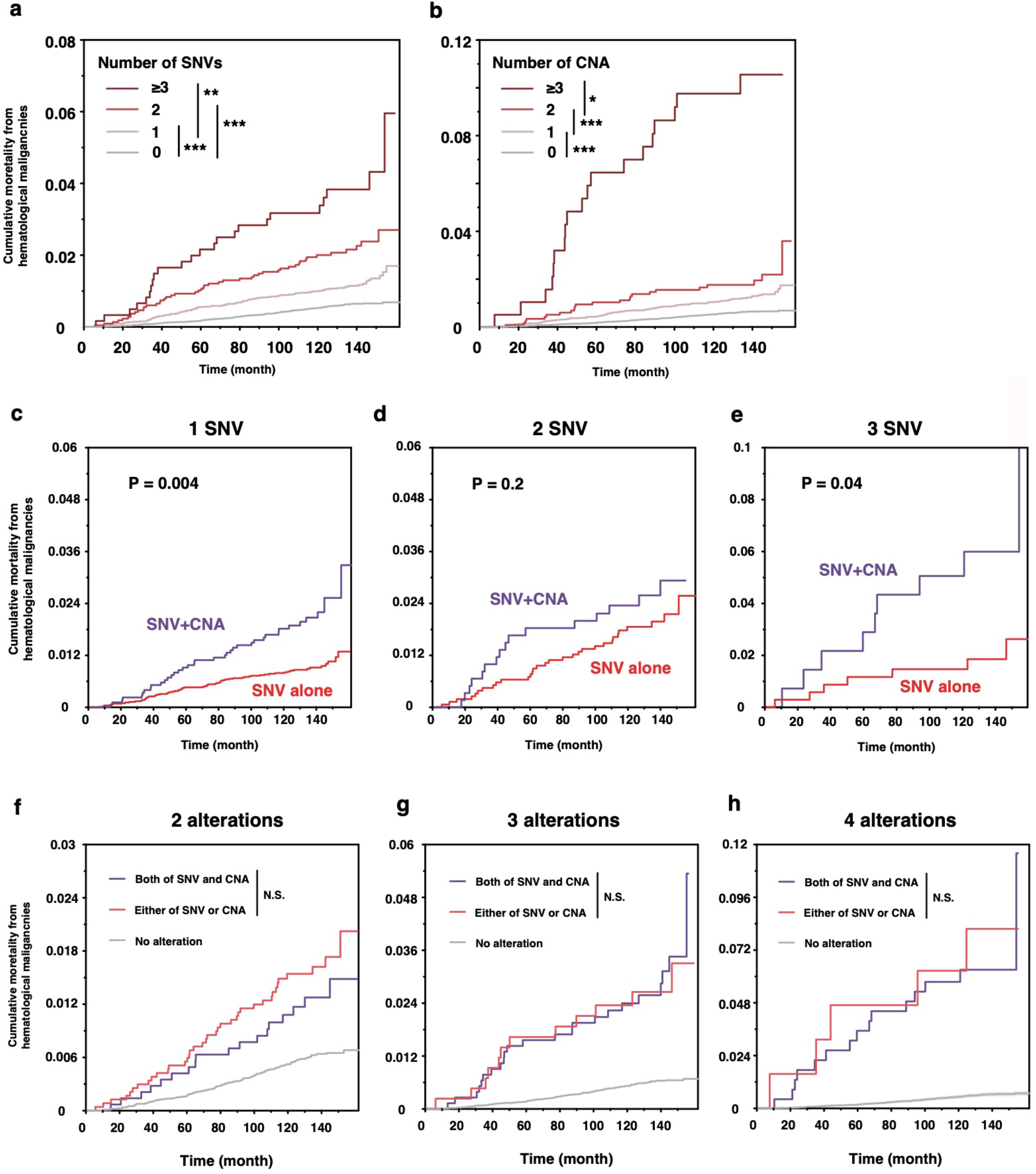
Impact of CH on mortality from HM stratified by number of alterations. a-b, Cumulative mortality from hematological malignancies in subjects with different number of SNVs/indels (a), or CNAs (b). c-e, Cumulative mortality from hematological malignancies in subjects with both SNVs/indels and CNAs or in those with SNVs/indels alone. Subjects with 1 (c), 2 (d), or ≤3 alterations are separately shown. f-h, Cumulative mortality from hematological malignancies in subjects with both SNV/indels and CNAs or in those with either of them. Subjects with 2 (f), 3 (g), or 4 alterations (h) are separately shown.

**Extended Data Fig. 9.**
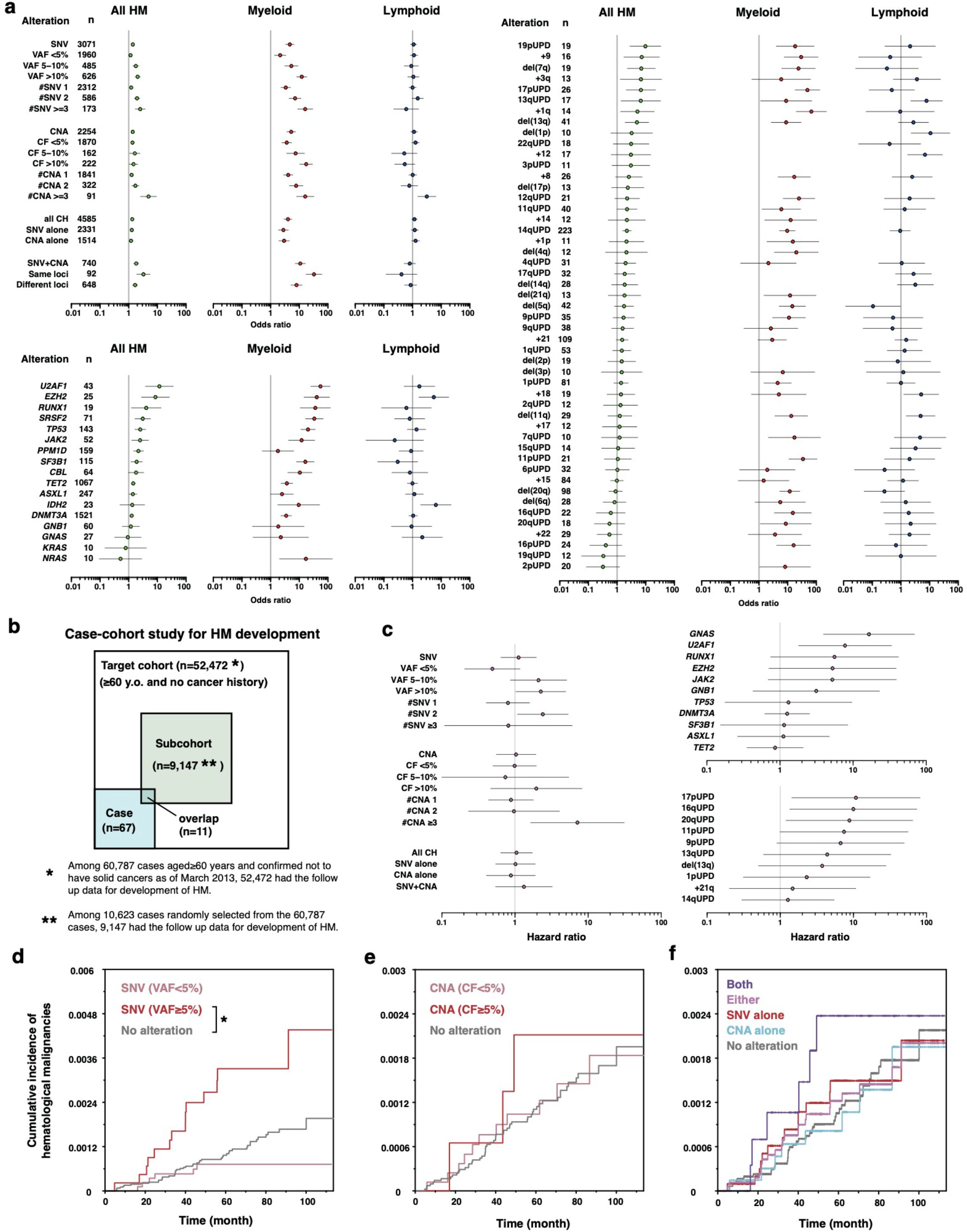
Interplay between SNVs/indels in *TP53* and CNAs. a, Odds ratios for the events (death and/or development) of hematological malignancies in case-control study (Extended Data Fig. 1a). b, Design of case-cohort study for development of hematological malignancies. c, Hazard rarios for development of hematological malignancies. d-f, Effect of SNVs/indels (d), CNAs (e), and combined SNVs/indels and CNAs (f) on the cumulative incidence of development of hematological malignancies. n, number of cases with the indicated alterations; SNV+CNA, coocurrence of both SNVs/indels and CNAs; #SNV, number of SNVs/indels; CF, cell fraction of CNAs; #CNA, number of CNAs.

**Extended Data Fig. 10.**
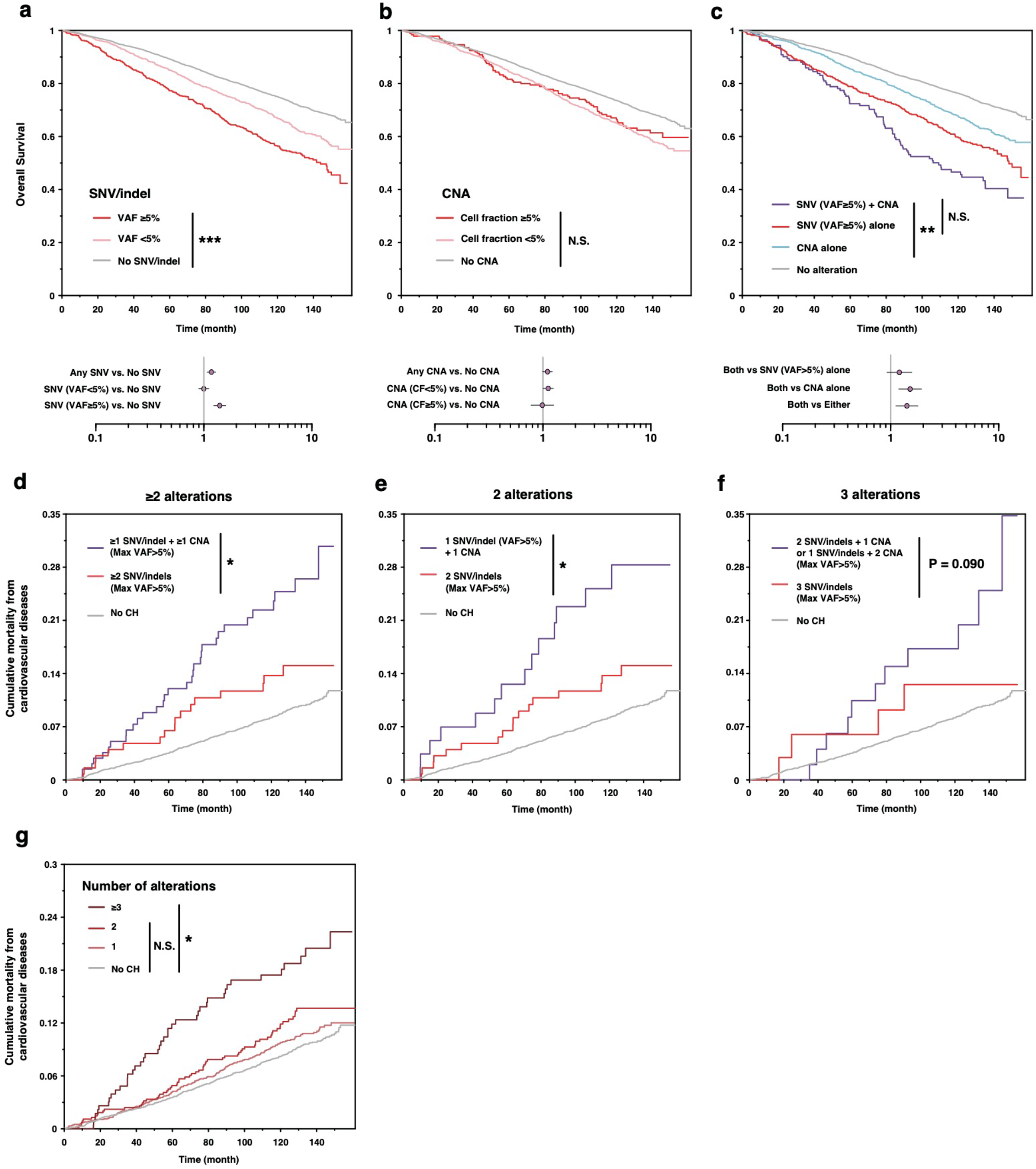
Interplay between SNVs/indels in *TP53* and CNAs. a-c,Effect of SNV/indels(a), CNAs(b), or combined SNV/indels and CNAs (c) on overall survivals. d-f, Cumulative mortality from cardiovascular diseases in subjects with SNVs/indels (Max VAF>5%) alone and those with both of SNV/indels (Max VAF>5%) and CNAs. Subject with 2≤ (d), 2 (e), and 3 (f) alterations are separately shown. g, Cumulative mortality from cardiovascular diseases in subjects with different number of CH-related alterations. N.S., not significant; *, P < 0.05; **, P < 0.001; ***, P < 0.0001.

