## Supplementary Tables for "Combined landscape of single-nucleotide variants and copy-number alterations in clonal hematopoiesis"

**Supplementary Table 1. Demographic summary of subjects.**

| Category | HM (+) (n = 672) | | HM (-) (n = 10,562) | |
|---|---|---|---|---|
| **Characteristics** | **No. of subjects (%)** | **Median (range)** | **No. of subjects (%)** | **Median (range)** |
| **Age at sampling (n = 11,234)** | | 71 (25-94) | | 70 (60-101) |
| <50 | 18 (2.7) | | 0 (0.0) | |
| 50-59 | 63 (9.4) | | 0 (0.0) | |
| 60-69 | 204 (30.4) | | 4,826 (45.7) | |
| 70-79 | 299 (44.5) | | 4,287 (40.6) | |
| 80-89 | 84 (12.5) | | 1,350 (12.8) | |
| 90-101 | 4 (0.6) | | 99 (0.9) | |
| **Gender (n = 11,234)** | | | | |
| Female | 234 (34.8) | | 4,933 (46.7) | |
| Male | 438 (65.2) | | 5,629 (53.3) | |
| **BMI (n = 10,532)** | | 22.9 (14-39.7) | | 23.1 (12-52.2) |
| <25 | 101 (15.9) | | 1,433 (14.5) | |
| ≥25, <30 | 514 (80.8) | | 8,115 (82.0) | |
| ≥30 | 21 (3.3) | | 348 (3.5) | |
| **History of smoking (n = 11,192)** | | | | |
| Current or past smoker | 387 (58.4) | | 4,997 (47.5) | |
| Non-smoker | 276 (41.6) | | 5,532 (52.5) | |
| **History of drinking (n = 11,158)** | | | | |
| Current or past drinker | 351 (53.4) | | 4,828 (46.0) | |
| Non-drinker | 306 (46.6) | | 5,673 (54.0) | |
| **Hypertension (n = 9,671)** | | | | |
| - | 387 (66.6) | | 5,440 (59.8) | |
| + | 162 (27.9) | | 2,817 (31.0) | |
| ++ | 29 (5.0) | | 678 (7.5) | |
| +++ | 3 (0.5) | | 155 (1.7) | |
| **Hyperlipidemia (n = 11,234)** | | | | |
| - | 179 (26.6) | | 3,895 (36.9) | |
| + | 493 (73.4) | | 6,667 (63.1) | |
| **Diabetes mellitus (n = 11,234)** | | | | |
| - | 171 (25.4) | | 3,177 (30.1) | |
| + | 501 (74.6) | | 7,385 (69.9) | |

HM (+/-): Subjects with/without event of hematological malignancies during follow-up periods.
Hypertension -: systolic blood pressures(sBP) < 140 and diastolic blood pressures(dBP) < 90, +: sBP≥140 or dBP≥90, ++: sBP≥160 or dBP≥100, +++: sBP≥180 or dBP≥110.

**Supplementary Table 2. Number of subjects with individual target diseases.**

| Disease group | Disease name | Number of subjects (%) | |
|---|---|---|---|
| | | **HM (+) (n = 672)** | **HM (-) (n = 10,562)** |
| Malignant tumors | Lung cancer | 27 (4.0) | 0 (0.0) |
| | Esophageal cancer | 13 (1.9) | 0 (0.0) |
| | Gastric cancer | 42 (6.3) | 0 (0.0) |
| | Colorectal cancer | 29 (4.3) | 0 (0.0) |
| | Liver cancer | 5 (0.7) | 0 (0.0) |
| | Pancreas cancer | 2 (0.3) | 0 (0.0) |
| | Gallbladder/Cholangiocarcinoma | 3 (0.4) | 0 (0.0) |
| | Prostate cancer | 45 (6.7) | 0 (0.0) |
| | Breast cancer | 20 (3.0) | 0 (0.0) |
| | Cervical cancer | 2 (0.3) | 0 (0.0) |
| | Uterine cancer | 4 (0.6) | 0 (0.0) |
| | Ovarian cancer | 1 (0.1) | 0 (0.0) |
| Cerebral diseases | Cerebral infarction | 90 (13.4) | 1740 (16.5) |
| | Cerebral aneurysm | 6 (0.9) | 208 (2.0) |
| | Epilepsy | 7 (1.0) | 89 (0.8) |
| Respiratory diseases | Bronchial asthma | 24 (3.6) | 494 (4.7) |
| | Pulmonary tuberculosis | **6** (0.9) | 29 (0.3) |
| | Chronic obstructive pulmonary disease | 17 (2.5) | 263 (2.5) |
| | Interstitial lung disease/Pulmonary fibrosis | 7 (1.0) | 68 (0.6) |
| Cardiovascular diseases | Myocardial infarction | 73 (10.9) | 1173 (11.1) |
| | Unstable angina | 31 (4.6) | 521 (4.9) |
| | Stable angina | 97 (14.4) | 1630 (15.4) |
| | Arrhythmia | 96 (14.3) | 1522 (14.4) |
| | Heart failure | 41 (6.1) | 909 (8.6) |
| | Peripheral arterial diseases | 19 (2.8) | 373 (3.5) |
| Liver diseases | Chronic hepatitis B | 4 (0.6) | 36 (0.3) |
| | Chronic hepatitis C | 29 (4.3) | 278 (2.6) |
| | Liver cirrhosis | 11 (1.6) | 83 (0.8) |
| Urologic diseases | Nephrotic syndrome | 4 (0.6) | 42 (0.4) |
| | Urolithiasis | 2 (0.3) | 263 (2.5) |
| Metabolic diseases | Osteoporosis | 44 (6.5) | 770 (7.3) |
| | Diabetes mellitus | 171 (25.4) | 3177 (30.1) |
| | Dyslipidemia | 179 (26.6) | 3895 (36.9) |
| Endocrine diseases | Graves' disease | 3 (0.4) | 74 (0.7) |
| Connective tissue diseases | Rheumatoid arthritis | 26 (3.9) | 292 (2.8) |
| Allergic diseases | Hay fever | 7 (1.0) | 176 (1.7) |
| Dermatologic diseases | Drug eruption | 1 (0.1) | 31 (0.3) |
| | Atopic dermatitis | 1 (0.1) | 13 (0.1) |
| | Keloid | 3 (0.4) | 19 (0.2) |
| Gynecologic diseases | Uterine fibroid | 2 (0.3) | 45 (0.4) |
| | Endometriosis | 0 (0.0) | 1 (0.0) |
| Pediatric diseases | Febrile seizure | 0 (0.0) | 0 (0.0) |
| Ophthalmologic diseases | Glaucoma | 21 (3.1) | 446 (4.2) |
| | Cataract | 87 (12.9) | 1991 (18.9) |
| Dental diseases | Periodontitis | 1 (0.1) | 131 (1.2) |
| Other | Amyotrophic lateral sclerosis | 0 (0.0) | 0 (0.0) |

HM (+/-): Subjects with/without event of hematological malignancies during follow-up periods.

**Supplementary Table 3. Summary of blood cell counts.**

| Blood cell count | HM(+) (n=672) | | HM(-) (n=10,562) | |
|---|---|---|---|---|
| | Median (range) | No. of subjects (%) | Median (range) | No. of subjects (%) |
| **White blood cell (/μL)** | 5450 (840-37500) | 551 | 5,900 (1,050-27,600 | 8,519 |
| Normal | | 507 (92.0) | | 8,092 (95.0) |
| ≥10000 | | 18 (3.2) | | 341 (4.0) |
| <3000 | | 26 (4.7) | | 86 (1.0) |
| **Hemoglobin (g/dL)** | 13.2 (4.8-17.7) | 573 | 13.5 (3.2-19.0) | 8,651 |
| Normal | | 517 (90.2) | | 7,814 (90.3) |
| ≥16.5 (male), 16 (female) | | 22 (3.8) | | 427 (4.9) |
| <10 | | 34 (5.9) | | 410 (4.7) |
| **Hematocrit (%)** | 39.7 (21.8-52.6) | 571 | 40.4 (15.5-69.4) | 8,645 |
| Normal | | 561 (98.2) | | 8,458 (97.8) |
| ≥50 | | 10 (1.8) | | 187 (2.2) |
| **Platelet ($10^4$/μL)** | 20.0 (1.1-131) | 511 | 21.0 (1.1-387) | 8,117 |
| Normal | | 477 (93.3) | | 7,920 (97.6) |
| ≥45 | | 7 (1.4) | | 54 (0.7) |
| <10 | | 27 (5.3) | | 143 (1.8) |

HM (+/-): Subjects with/without event of hematological malignancies during follow-up periods.

**Supplementary Table 4. Antibodies for cell sorting.**

| Antibody | Catalog number | Manufacturer | Clone |
|---|---|---|---|
| FITC anti-human CD19 | 560994 | BD Bioscience | HIB19 |
| PE anti-human CD3 | 552127 | BD Bioscience | SP34-2 |
| APC anti-human CD235a | 561775 | BD Bioscience | HIR2 |
| PE-Cy7 anti-human CD34 | 343516 | Biolegend | 581 |
| BV421 anti-human CD33 | 744761 | BD Bioscience | P67.6 |
| BV421 anti-human CD13 | 744862 | BD Bioscience | L138 |

**Supplementary Table 5. Primer sequences for detection of allele imbalances in the regions of del(13q).**

| SNP ID | Status | SNP position | Forward primer sequence | Reverse primer sequence |
|---|---|---|---|---|
| rs731779 | Deleted | chr13:47452038 | AAGCGGCCGCAAAGCAGGGCAAGTACCTCA | AAGCGGCCGCTGAGTGTCTCTCTTGCCCCA |
| rs1350457 | Deleted | chr13:54355150 | AAGCGGCCGCGGTAAGAATACAAACCTGGAAAAAGTG | AAGCGGCCGCCCTTGGACCCGCTTCACTC |
| rs341506 | Deleted | chr13:60420314 | AAGCGGCCGCACACAGGCTTTCCTCCAAGT | AAGCGGCCGCTGTGTAAGAGTGAGTGTGGCA |
| rs359362 | Deleted | chr13:65239972 | AAGCGGCCGCTTGGTCAAATGGCACCCCTT | AAGCGGCCGCCAATTAGATTTGGAATTTGCTTGTGA |
| rs4773419 | Intact | chr13:112311079 | AAGCGGCCGCAAGAAAGGCAGGTCCAAGGG | AAGCGGCCGCGTGTTGACAAAGCCGGTTGG |

**Supplementary Table 6. Probes used for ddPCR.**

| Gene | Amino acid substitution | BioRad Assay ID |
|---|---|---|
| *TET2* | p.A1153V | dHsaMDS869039740 |
| *JAK2* | p.V617F | dHsaMDS488977115 |
| *TP53* | p.R175H | dHsaMDV2010105 |
| *TP53* | p.Y220C | dHsaMDV2510536 |
| *TP53* | p.R248Q | dHsaMDV2010127 |
| *TP53* | p.R248W | dHsaMDV2010107 |
| *TP53* | p.R273H | dHsaMDV2010109 |
| *TP53* | p.R273C | dHsaMDV2510538 |
