## Supplementary Figures for "Combined landscape of single-nucleotide variants and copy-number alterations in clonal hematopoiesis"

### Supplementary Fig. 1

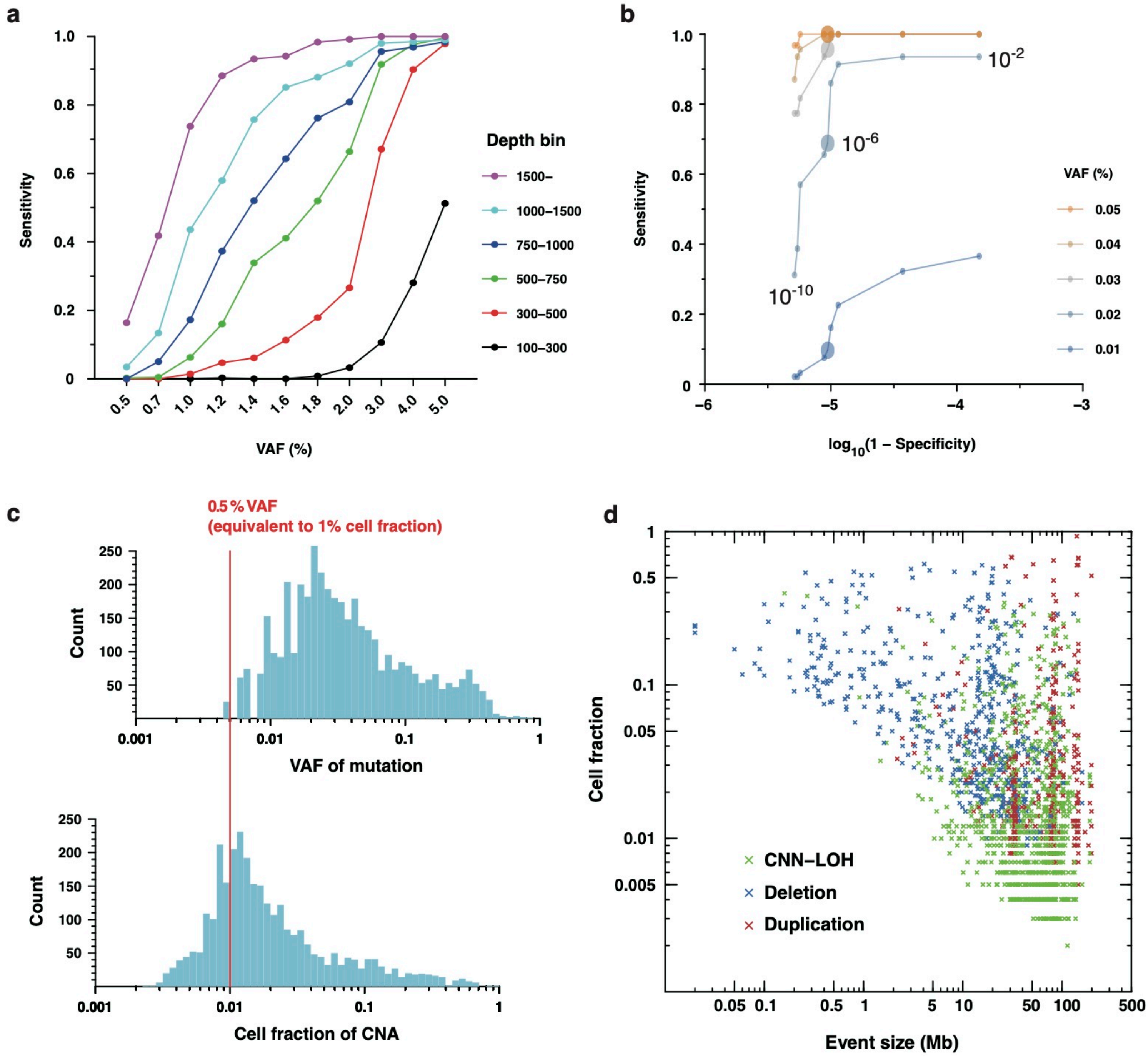


**Supplementary Fig. 1 I Distributions of clone sizes and performance evaluation.**

a, Sensitivities to detect SNVs simulated for different VAFs and sequencing depths. The horizontal axis represents target VAF of simulated SNVs. The vertical axis represents sensitivity, which was calculated as fractions of detected SNVs out of all simulated ones. b, Receiver operating characteristic (ROC) curves for detection of SNVs/indels, illustrating sensitivity on the vertical axis and $\log_{10}$(1 - specificity) on the horizontal axis. In this panel, we show sensitivity assuming sequencing depth is within x700-x900, which largely represents for the mean coverage in this study (x800). Dots represent variable cutoffs on beta-binominal P values ($10^{-2}$ to $10^{-10}$) (Online Method). Large dots represent a cutoff of $10^{-6}$, which we adopted in the actual SNV call. c, Histograms of VAFs of SNVs/indels (top), and cell fractions of CNAs (bottom). The red vertical line indicates 0.5 % in VAFs, which is equivalent to 1% in cell fractions. It was impossible to precisely calculate cell fractions for unclassifiable CNAs. Instead, we calculated upper limits of cell fractions by assuming they were duplication. d, Distribution of detected CNAs with cell fractions on the vertical axis and event sizes on the horizontal axis. Unclassifiable CNAs are abbreviated from panel (d).

### Supplmentary Fig. 2

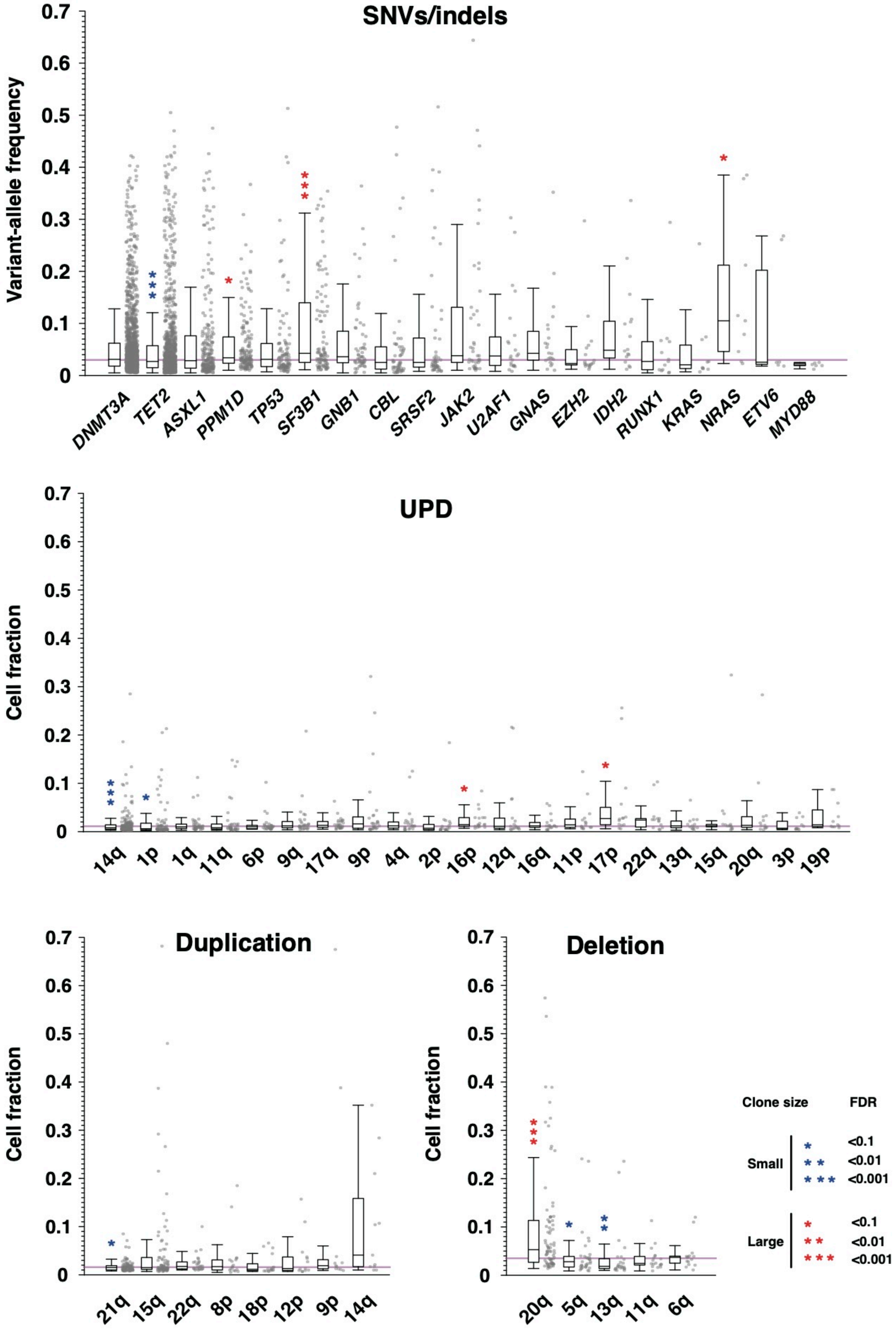


**Supplementary Fig. 2 | Clone size of individual CH-related alterations**

VAF/clone size of individual CH-related alterations are compared within each category (SNVs/indels, UPDs, duplications, and deletions). FDR are calculated in comparison with clone size of all other alterations by wilcoxon rans-sum test. Purple horizontal lines indicate median clone size within each category.

### Supplementary Fig. 3

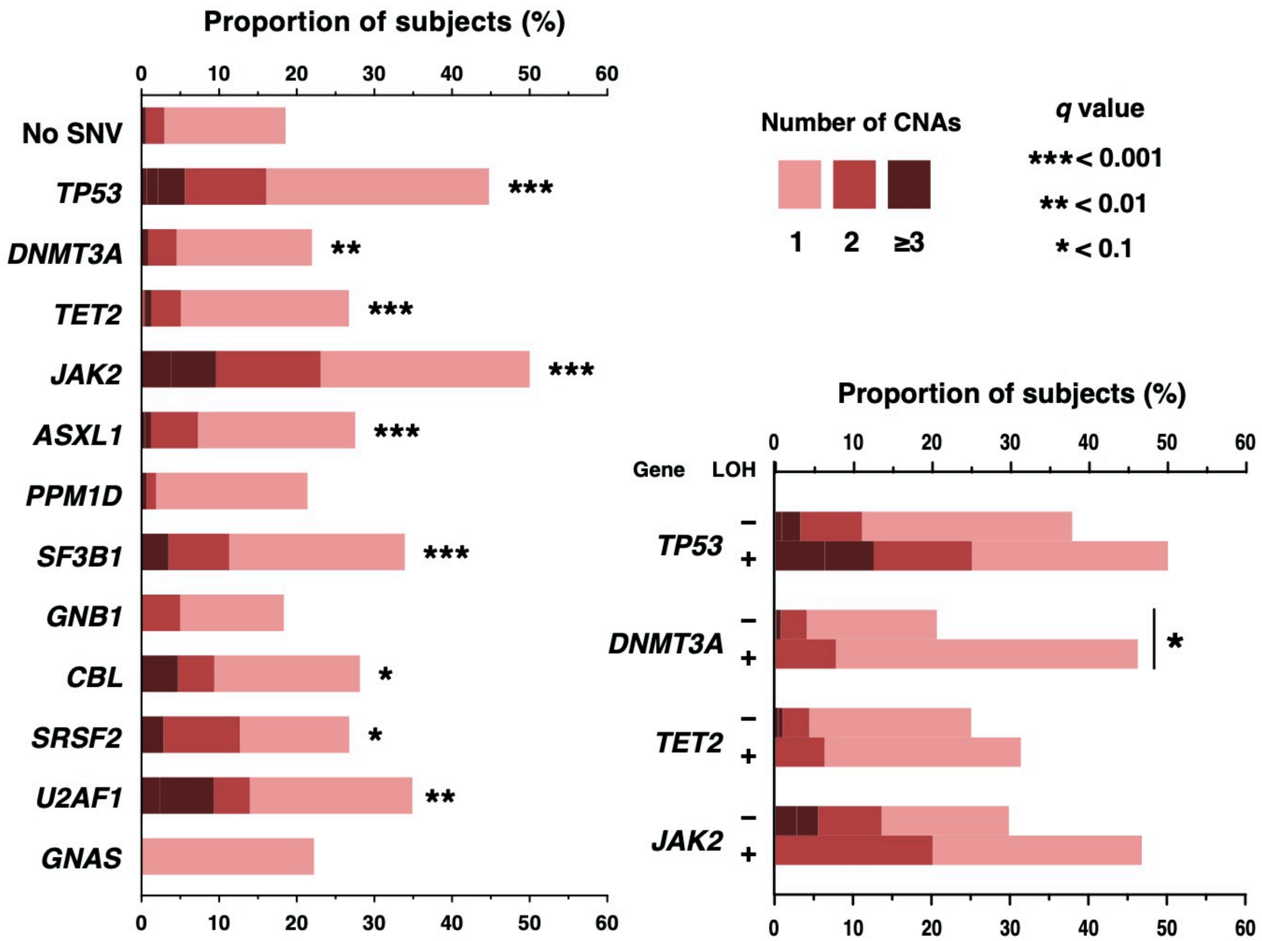


**Supplementary Fig. 3 | Relationships of SNVs/indels and number of cooccurring CNAs.**
Proportions of subjects with CNAs within those who harbor SNVs/indels in the indivated genes (left) and comparison of the number of cooccurring CNAs between subjects with SNVs/indels with or without LOH in *TP53*, *DNMT3A*, *TET2*, and *JAK2* (right). The proportions of subjects with 1, 2, and ≥3 CNAs are depicted by different colors. In the left, the numbers of CNAs are compared with subjects without SNVs/indels (labeled as “No SNV” ) by Wilcoxon test. In LOH(+) groups in the right panel, we did not counted CNAs responsible for the LOH in the number of coocurring CNAs. Significantly larger numbers of CNAs are indicated by

### Supplementary Fig. 4

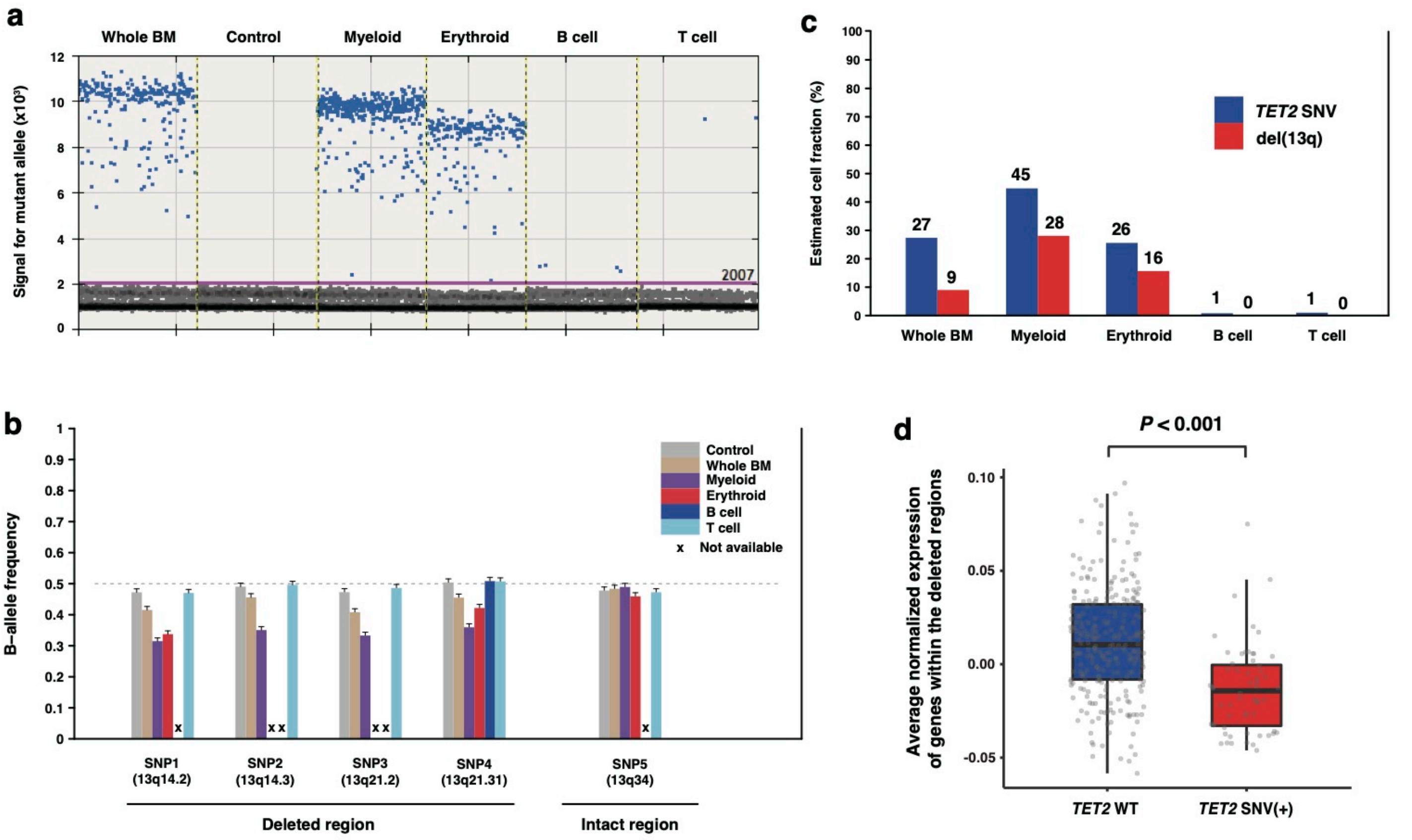


**Supplementary Fig. 4 | Analysis of a representative case with a SNV in *TET2* and del(13q).**

a, Results of ddPCR for A1153V substitution in *TET2* performed on DNA samples extracted from whole bone marrow cells, myeloid cells (CD13/33+), erythroid cell (CD235a+), B cells (CD19+), and T cells (CD3+). Sample for negative control is taken from a CH-negative subject. b, B-allele frequencies (BAF) for 4 heterozygous SNPs (Supplementary Table 6) within del(13q) and one in an intact region. Beucase of small amounts of DNA, BAF for SNP1, 2, 3, and 5 were not available for B cell, and that for SNP 2 and 3 were not available for erythroid. Error bars indicate upper limits of 95% confidence intervals. c, Cell fractions of the SNV in *TET2* and del (13q) in each fraction. Cell fraction for the *TET2* SNV was calculated as 2 × VAF. That for del(13q) were calculated on the basis of allelic imbalance observed at SNP4 in panel (b). d, Results of single-cell gene expression analysis and SNV detection in Fluidigm C1 platform. Average normalized expression of genes within del(13q), which can be a surrogation for DNA copy-number of the deleted region, are plotted for each cell with or without A1153V substition in *TET2*. The box plot indicates the median, first and third quartiles (Q1 and Q3) and whiskers extend to the furthest value between Q1 – 1.5×the interquartile range (IQR) and Q3 + 1.5×IQR.

### Supplementary Fig. 5

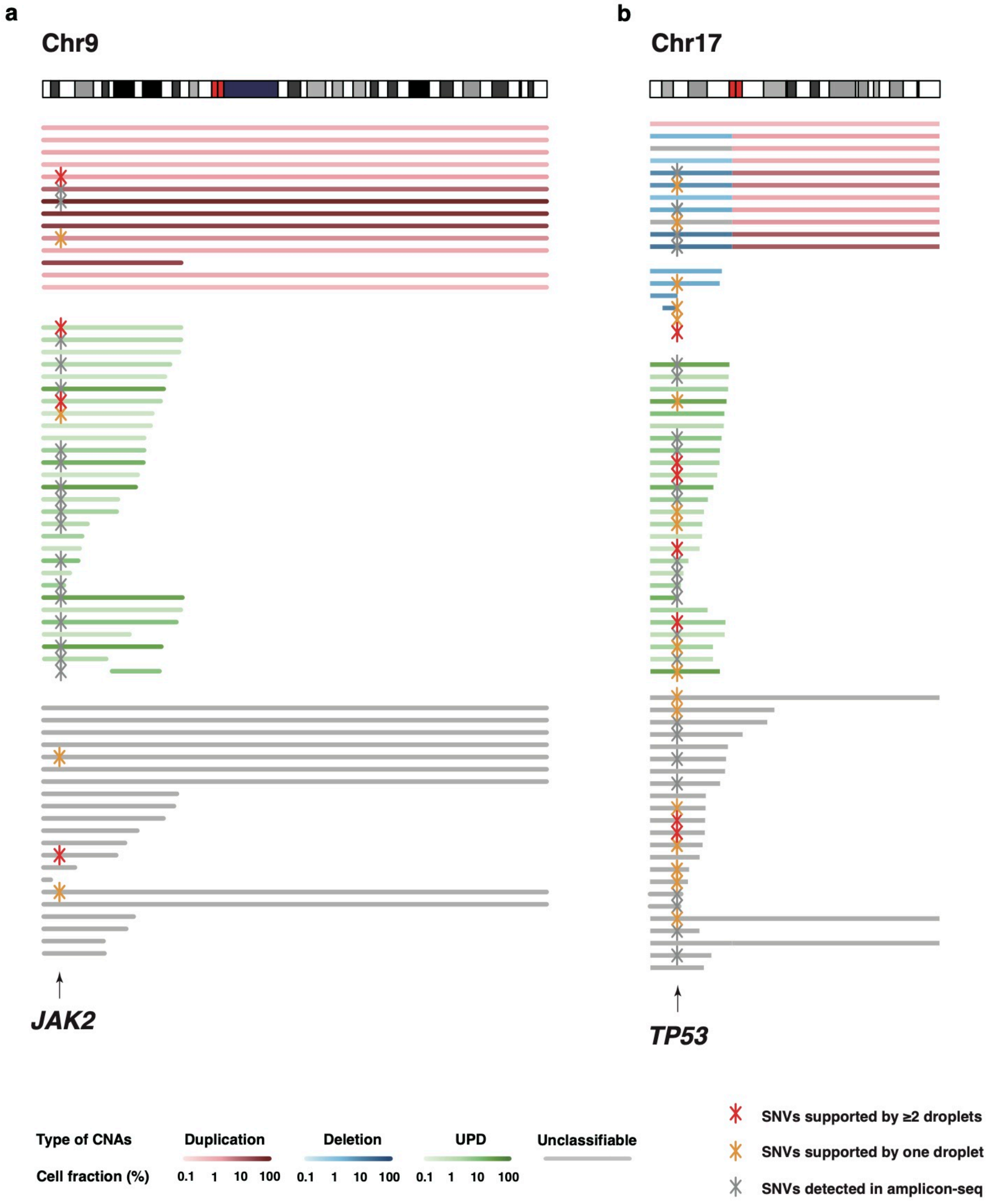


**Supplementary Fig. 5 | ddPCR for mutational hotspots in *JAK2* and *TP53*.**

a-b, Hotspot SNVs newly detected in ddPCR are illustrated by red or orange asterisks. We tested V617F in *JAK2* and R175H, Y220C, R248Q/W, R273C/H in *TP53*. SNVs supported by multiple droplets are shown in red, those suppported by single in orange, and those already detected in targeted sequencing in gray.

### Supplementary Fig. 6

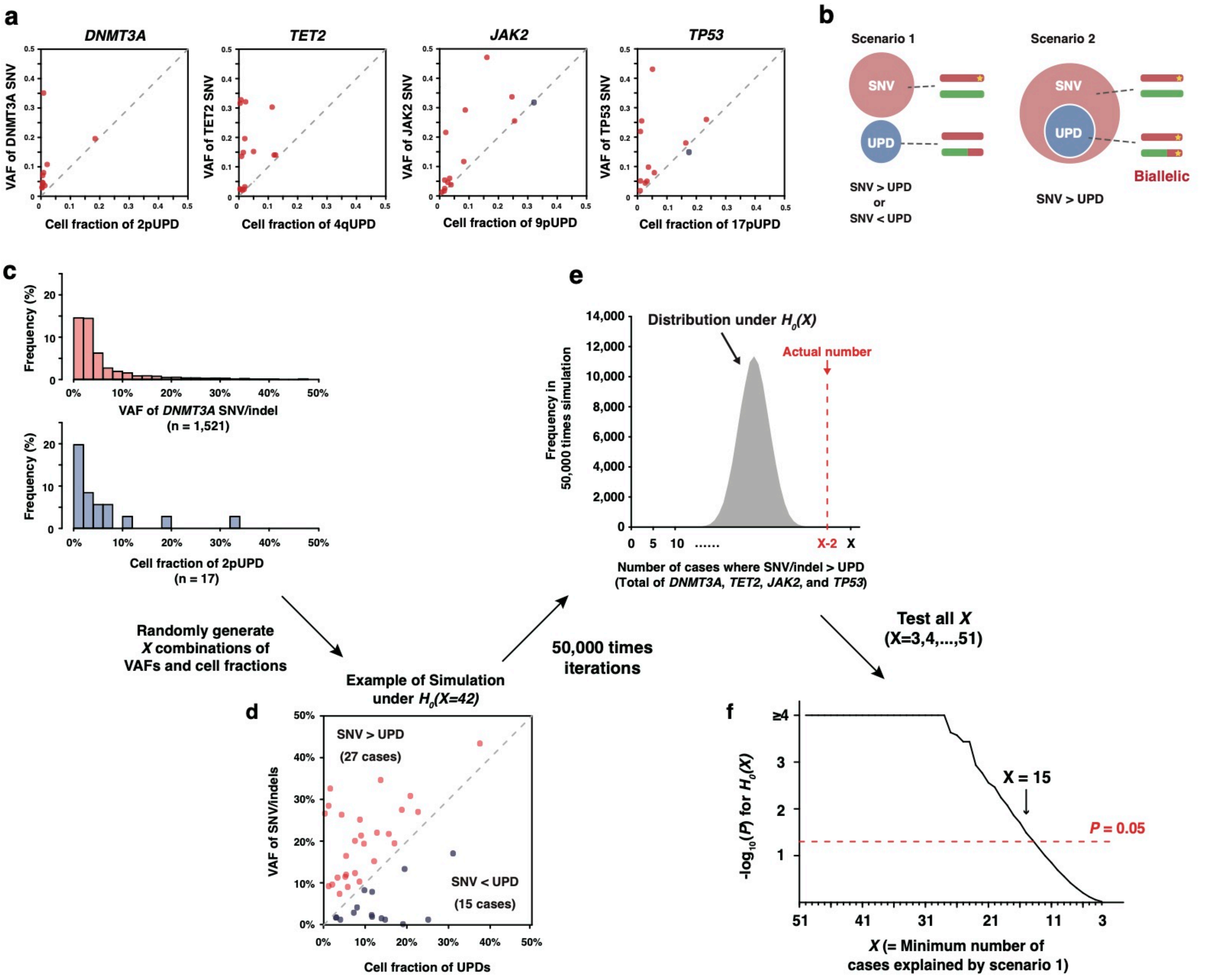


**Supplementary Fig. 6 | Simulation for VAF/cell fractions of coocurring SNVs/indels and UPDs.**

a, Comparison of VAFs and cell fractions of SNV/indels and UPDs involving the same genes: *DNMT3A* (n=10, a), *TET2* (n=15, b), *JAK2* (n=14, c), and *TP53* (n=12, d). Compared with cell fractions of UPDs, VAF of SNVs/indels were larger in 49 of the 51 (red) and smaller in the remaining 2 cases (blue). b, Two possible scenarios underlying the observations in (a). In scenario 1, SNVs/indels and UPDs exist in discrete cells. In that case, VAF of SNVs/indels can be larger or smaller than cell factions of UPDs. In scenario 2, UPD subclonally exists within the clone carrying SNVs/indels, causing biallelic alterations. The 51 observations in (a) can be regarded as a mixture of the two scenarios. In the following simulation, we put a null hypothesis $H_0(X)$, that at least X of the 51 cases in (a) are explained by scenario 1. c, Histgrams of VAFs of *DNMT3A* SNVs/indels and cell fractions of 2pUPDs in the entire cohort. In the simulation, we randomly sample *X* combinations of the VAF of SNVs/indels and cell fractions of UPDs from these distributions. Only distributions of *DNMT3A*/2pUPD are shown, but we also sample VAFs and cell fractions from the distributions of *TET2*/4qUPD, *JAK2*/9pUPD and *TP53*/17pUPD as well (not shown). d, An example of simulated combinations of VAF and cell fractions. Here, we supposed *X*=42 and simulated VAFs of SNVs/indels were bigger than cell fractions of UPDs in 27 cases. e, Iterating the procedures illustrated in (c) and (d), we obtained a null distribution of the number of cases in which VAFs of SNVs/indels were larger than cell fractions of UPDs (shown in gray). Comparing the null distribution with the actually observed number, *X*-2 (shown in red), we calculated P value for $H_0(x)$ (x=3,...,51) and looked for the minimum *X* with *P*<0.05. f, *P* values for $H_0(x)$ (x=3,...,51) are shown. The minimum *X* with *P* < 0.05 was 15, which suggested scenario 1 can explain less than 15 cases out of the 49 cases in which VAFs of SNVs/indels were larger than cell fractions of UPDs. Thus, the remaining 34 cases should be explained by scenario 2.

### Supplementary Fig. 7

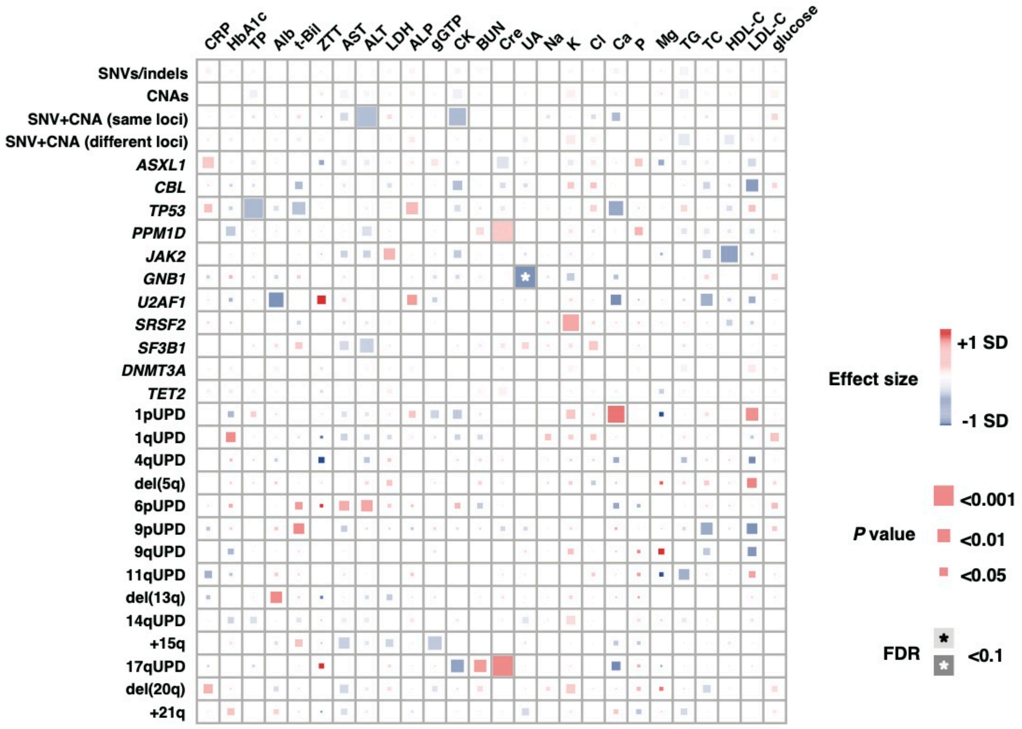


**Supplementary Fig. 7 | Association of CH wtih blood test values.**

Positive or negative correlation between CH-related alterations and blood test values are illustrated in red or blue rectangles, respectively. CRP, C-reactive protein; HbA1c, Hemoglobin A1c; TP, total protain; Alb, albumine; t-Bil, total bilirubin; ZTT, zinc sulfate turbidity test; AST, aspartate aminotransferase; ALT, alanine aminotransferase; LDH, lactate dehydrogenase; ALP, alkaline phosphatase; gGTP, gamma-glutamyltransferase; CK, creatinine kinase; BUN, blood urea nitrogen; Cre, creatinine; UA, uric acid; Na, sodium ion; K, potassium ion; Cl, chloride ion; Ca, calcium ion; P, phosphate ion; Mg, magnesium ion; TG, triglycerol; TC, total choresterol; HDL-C, high-density lipoprotein cholesterol; LDL-C, low-density lipoprotein cholesterol.

### Supplementary Fig. 8

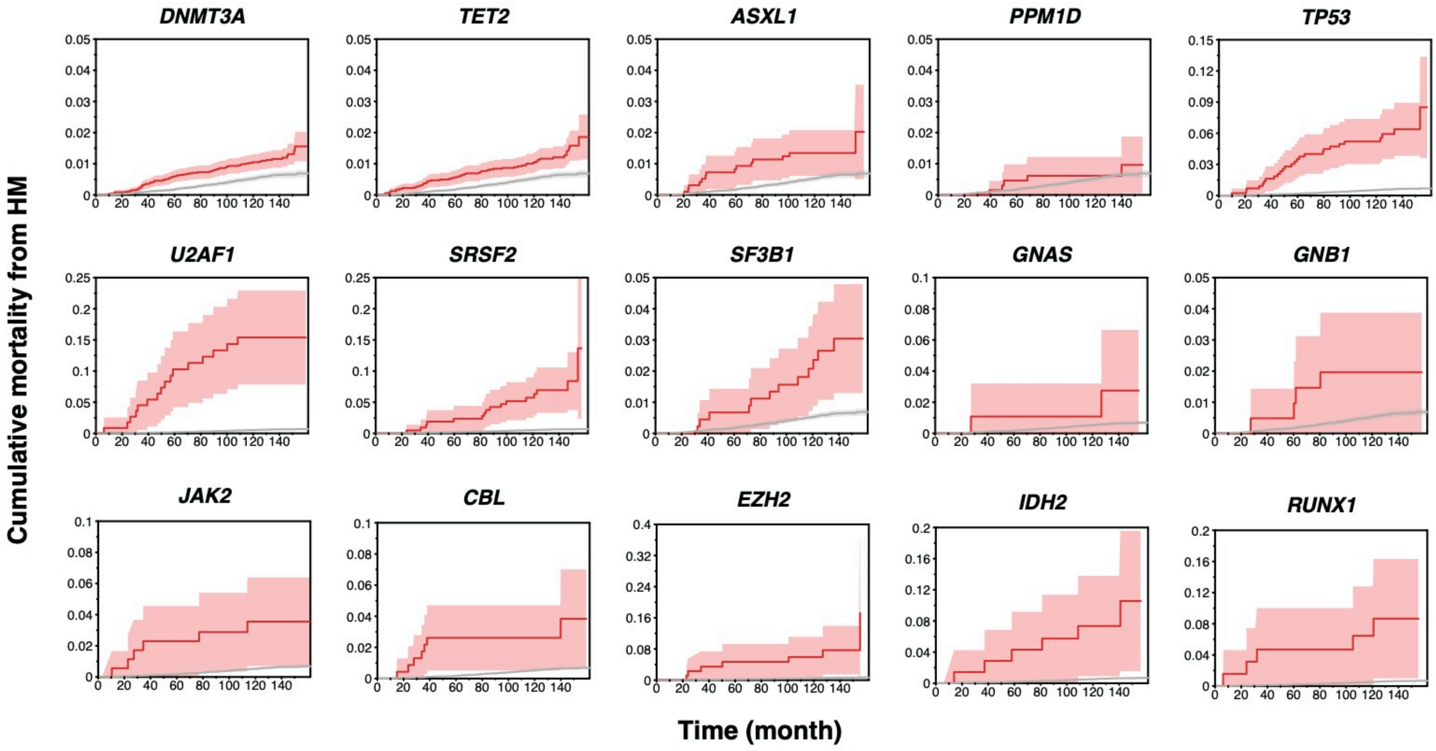


**Supplementary Fig. 8 | Cumulative mortality from HM in subjects with individual SNVs/indels.**

Cumulative mortality from HM in subjects with SNVs/indels in the indicated genes. For comparison, cumulative mortality from HM in subjects without any alteration is also shown in gray. Colored bands indicate 95% confidence intervals.

### Supplementary Fig. 9

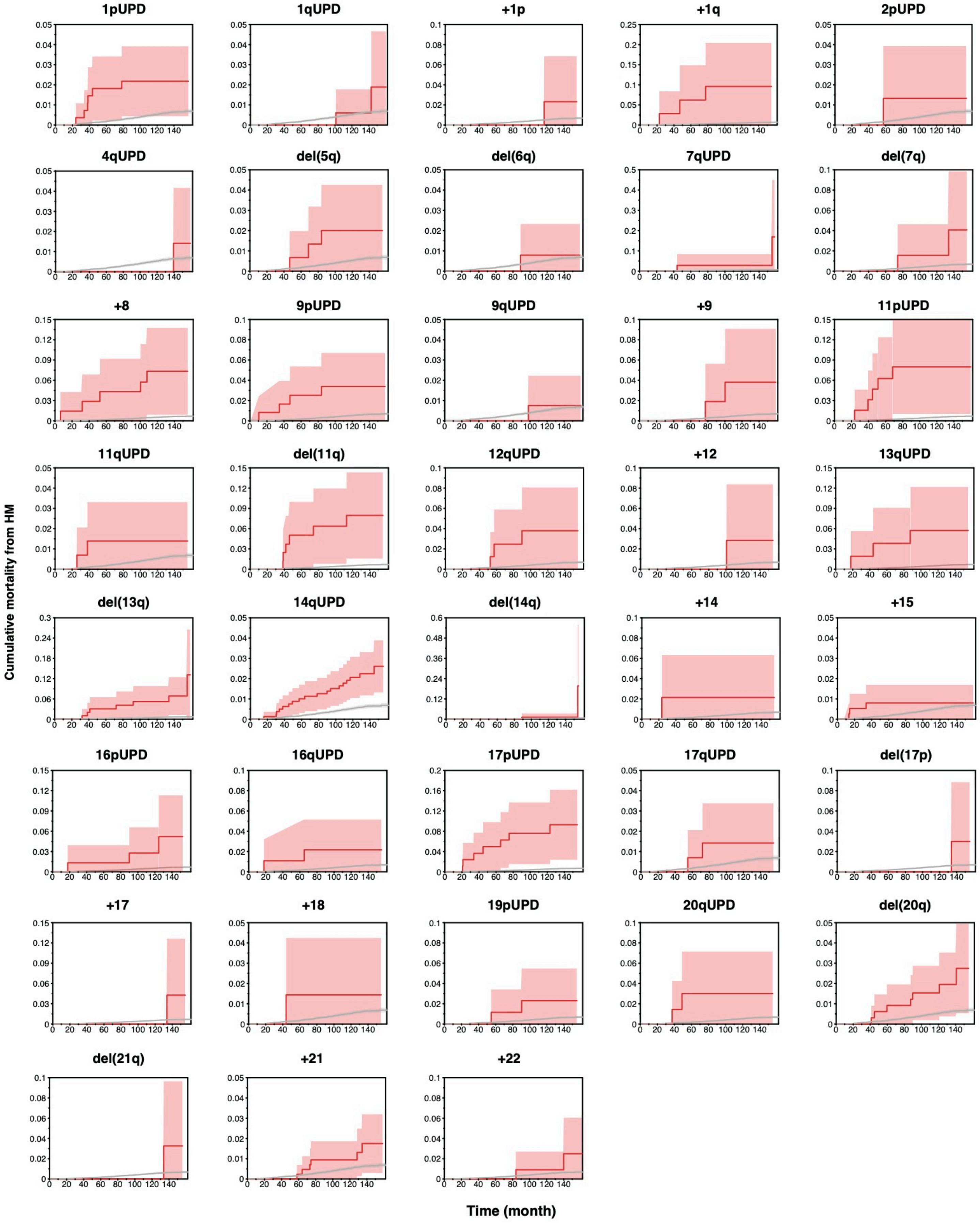


**Extended Data Fig. 9 | Cumulative mortality from HM in subjects with individual CNAs.**

Cumulative mortality from HM in subjects with the indicated CNAs. For comparison, cumulative mortality from HM in subjects without any alteration is also shown in gray. Colored bands indicate 95% confidence intervals.
